# Stitching flexible electronics into the brain

**DOI:** 10.1101/2023.04.20.537740

**Authors:** Jung Min Lee, Dingchang Lin, Young-Woo Pyo, Ha-Reem Kim, Hong-Gyu Park, Charles M. Lieber

## Abstract

Understanding complex neuronal networks requires monitoring long-term neuronal activity in various regions of the brain. Significant progress has been made in multi-site implantations of well-designed probes, such as multi-site implantation of Si-based and polymer-based probes. However, these multi-probe strategies have been limited by the sizes and weights of interfaces to the multiple probes and the inability to track the activity of the same neurons and changes in neuronal activity over longer time periods. Here, we report a long single flexible probe that can be implanted by stitching into multiple regions of the mouse brain and subsequently transmit chronically-stable neuronal signals from the multiple sites via a single low-mass interface. We implanted the probe at four different sites using a glass capillary needle or two sites using an ultrathin metal needle. In-vitro tests in brain-mimicking hydrogel showed that multi-site probe implantations achieved a high connection yield of >86%. In-vivo histological images at each site of probes, implanted by stitching using either glass capillary or ultrathin metal insertion needles exhibit seamless tissue-probe interfaces with negligible chronic immune response. In addition, electrophysiology studies demonstrated the ability to track single neuron activities at every injection site with chronic stability over at least one month. Notably, the measured spike amplitudes and signal-to-noise ratios at different implantation sites showed no statistically significant differences. Multi-site stitching implantation of flexible electronics in the brain opens up new opportunities for both fundamental neuroscience research and electrotherapeutic applications.

## 1. Introduction

Implantable neural probes^1-6^ continue to play an important role in neuroscience research, and represent important and/or possible therapeutic options for neurological diseases and disorders such as Parkinson’s disease and major depressive disorder.^7-9^ To understand normal and aberrant processing in neural networks, neural probes should ideally be able to monitor many neurons with high spatial and temporal resolution,^10-12^ to access multiple regions in the brain encompassing a network of interest, and ultimately to be capable of monitoring awake, freely behaving subjects. Several electronic probe strategies have been pursued for recording neural signals simultaneously from distinct brain regions necessary for single neuron-resolved network studies. For example, multiple implanted polymer-based neural probes have been capable of chronically detecting brain signals in distinct brain regions, although the quantity and complexity of input/output (I/O) interfaces increased along with the number of implanted probes.^13-15^ Multiple single-shank and multi-shank array Si-based probes also have been used to probe distinct brain regions,^10-12,16,17^ but their capacity to track the same neurons over months is constrained by chronic gliosis and instability caused by mechanical and structural mismatches with the brain tissue.^18,19^ The Si- and polymer-based probes also typically have increasing large mass I/O amplifier interfaces, which have either wired^13,16^ or wireless^20,21^ connections to recording electronics, when using multiple probes. In addition, multi-shank array Si-based probes can only target very specific brain regions without being tunable due to the limitation of fixed design and fabrication.^16,17^

## 2. Results

In this work, we asked whether it would be possible to address these issues by using a single flexible probe that is implanted into two or four sites of the mouse brain, including sites spanning the left/right hemispheres, and recording detected signals from the distinct regions via a single interface. This strategy is illustrated schematically with a single flexible probe is implanted into four different regions across both left and right hemispheres of the mouse brain (Figure 1a). We focus on probes with a mesh-like design that have previously shown tissue-like mechanics and structure for chronic biocompatibility in single-site implantations by syringe injection.^22-29^ Uniquely here, we ask whether this implantation strategy can be exploited to stitch, much like when sewing with a needle and thread, the single long probe into the hippocampal regions of both hemispheres of the mouse brain without being constrained to a specific set of pre-determined locations. In so doing, the multi-site could be accessed using a single interface of low-profile low-mass (∼1 g) headstage including the flat flexible cables (FFC), the customized printed circuit board (PCB) and the amplifier is necessary for multiplexed I/O interfaces with recording instruments (Figure 1a).

**Figure 1.**
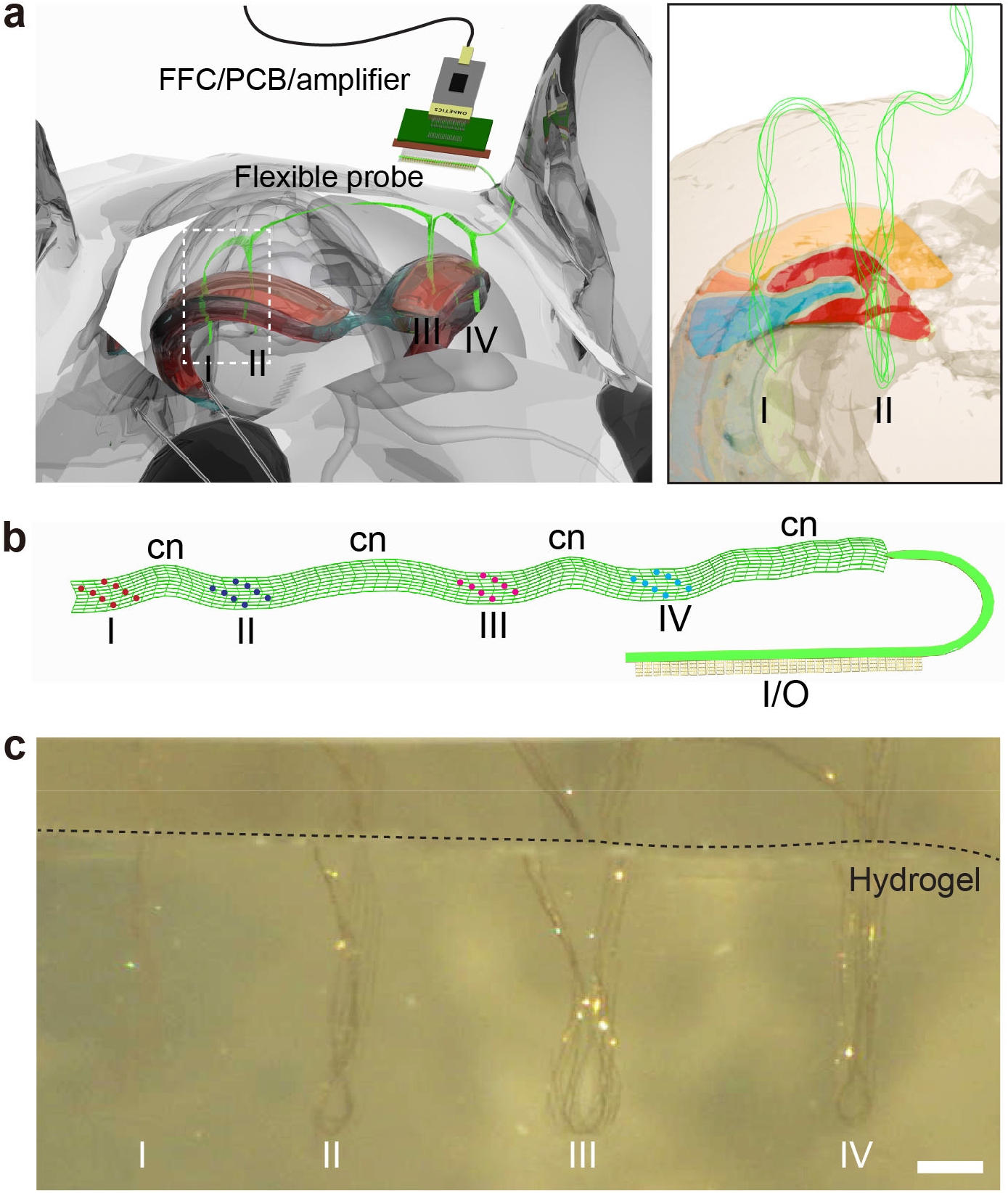
Overall concept of stitching flexible electronics using glass needle. a) Schematic of four-site implantations (I, II, III, and IV) of a single flexible probe (green) across the mouse brain using a glass needle with a single headstage/interface. Right, magnified schematic of (a) flexible probe implanted in different brain regions (I and II). b) Design of a flexible probe design for four-site implantations. The single flexible probe consists of four probing regions (I, II, III, and IV), as well as their connectors (cn) and an I/O (input/output) pad. There are eight recording electrodes in each probing region. c) Optical microscope image of a single flexible probe implanted into 0.5% agarose hydrogel from four different regions (I to IV). Scale bar, 3 mm.

To faciliate four-site implantations with a stable probe/neuron interface and high spatial resolution, the following points were considered when designing the flexible probe: (1) The bending stiffness should be low enough to be comparable to tissues, allowing the seamless integration between the tissue and the probe as well as stable chronic recording of the same neurons over a few months; (2) The width of the probe should be determined by the inner diameter of the syringe being loaded, allowing for smooth injection of the probe without crumpling; (3) The distribution of recording electrodes in the probe should be determined by the target region and depth of the brain, to precisely position the electrodes in desired areas. Points (1) and (2) have been described previously for reported mesh electronics probes.^27-29^ Point (3) is addressed in the design phase specific to given experiments such that the groups of electrodes, 2 or 4 for two- or four-site studies, can be targeted to the different brain regions with sufficient long connection regions between the electrode groups. Figure 1b shows a schematic flexible mesh probe for four-site implantations that meets these requirements. The total length and width of the flexible probe were 30 mm and 1 mm, respectively, while the thickness of the probe and the angle between the longitudinal and transverse ribbons were the same as in our previous design (Figure S1a).^27-29^ In addition, the platinum (Pt) recording electrodes were distributed across the four distint regions (I; red, II; blue, III; pink, and IV; cayn in Figure 1b). The Pt recording electrodes of each region and I/O pads were electrically connected by connectors (cn).

To implant a single flexible probe into the brain multiple times, we developed a stitching process, which included the following steps (Figure 1c, Figure S2, and Movie S1). First, the flexible probe is loaded with sterile saline in a capillary needle and stereotaxically implanted into the left hippocampus (Figure S2, i). Second, the needle is withdrawn and moved to the next target position using the motorized stereotaxic frame (Figure S2, ii). The first and second processes are then repeated in the subsequent insertion sites, which include target areas in the right hippocampus (Figure S2, iii and iv).

To confirm whether the flexible probe was not damaged or broken even after multiple implantations, the impedance of the electrodes was measured in three parallel devices after four-site implantations into 0.5% agarose hydrogel (Figure 1c). We chose hydrogel because its mechanical properties are similar to those of brain tissues.^30,31^ The impedance measurement showed that a high connection yield of ca. 89% was successfully achieved (Figure S3): 85 out of 96 channels were connected to the FFC in the in-vitro condition, with impedance values less than 1 MΩ. This in-vitro test of four-site implantations of a single flexible probe using a needle highlights several key points. First, although the new probe design was ten times longer than our previous one,^22-29^ we smoothly injected the probe at a rate of 10−20 mL/h in the first implantation. Second, the probe was implanted precisely and less crumpled because the mesh was implanted without the aid of liquid flow after the first implantation. Last, multi-site implantations in hydrogel also demonstrated a high connection yield even with the probe structure ten times longer than previous ones.^22-29^

After the successful demonstration of in-vitro stitching tests of four-site implantations, we asked whether in-vivo four-site implantations show a stable and seamless probe electronics/brain tissue interface and allow for long-term single-neuron mapping.^23,26^ To address these questions and investigate time-dependent histology, we implanted a single flexible probe into the hippocampal regions of both hemispheres in the mouse brain at four different sites (Figure 2a, i-iv). Following the initial implantation at site-I, optical imaging of the brain surface shows that the mesh probe connecting implantation sites is on the surface of the brain (blue arrows in Figure 2a, iv; Methods). In this experiment, we used transgenic mouse lines expressing either yellow fluorescent proteins (YFPs) in neurons (Thy1-YFP-H) or green fluorescent proteins (GFPs) in astrocytes (GFAP-GFP). Figure 2b shows a representative three-dimensional (3D) reconstructed image of the probe and neurons in both hemispheres at 2 weeks post-injection.^23^ A single probe was successfully implanted into four different regions of the brain of a mouse (Figure 2b, I-IV). The probe from these regions showed linearly elongated conformations without crumpling, interpenetrating through the cortex (CTX), hippocampus (HIP) and dentate gyrus (DG). Higher-resolution images of the hippocampus were also taken for quantitative analysis of probe/neurons or probe/astrocytes interfaces after four-site implantations, using several independent samples at 2 and 4 weeks post-injection in both Thy1-YFP-H and GFAP-GFP transgenic mice (N = 2 each, total = 8) (Figures S4 and S5). To assess the probe/brain interface, we quantified the normalized fluorescence intensity of neurons as a function of the distance from the probe surface in the high-resolution 3D images. For further clarification of the fluorescence intensity of neurons and astrocytes, we plotted the average intensity of neurons and astrocytes inside and outside of the probe at 2 and 4 weeks post-injection (Figures S4c and S5c). These results show a relatively uniform tissue population with no discernible depletion of neurons or accumulation of astrocytes near the probe at both time points. Thus, there is no evidence that the four-site implantations cause a chronic immune response or adversely affect the probe-tissue interface over time.

**Figure 2.**
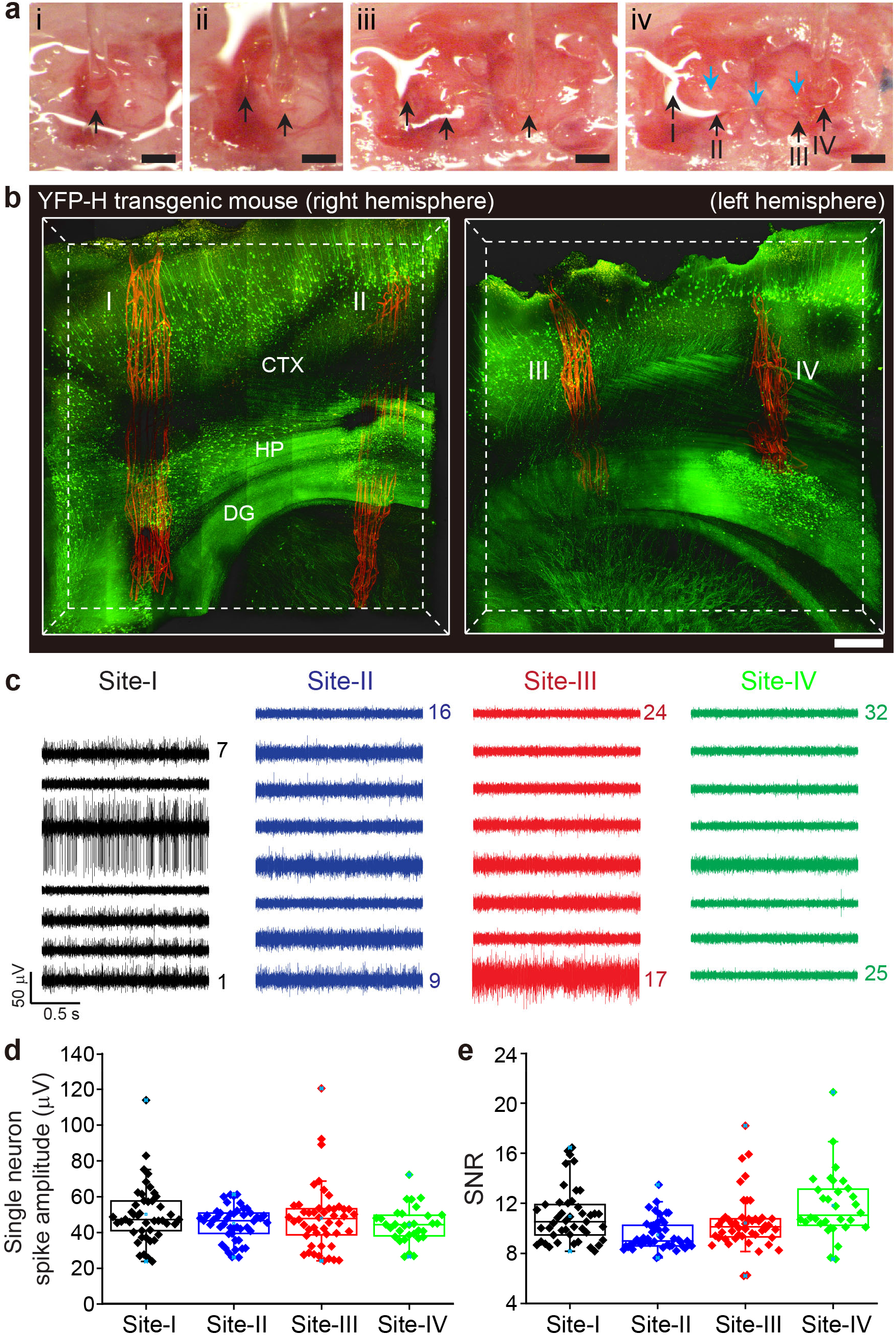
Four-site implantation of flexible electronics using a glass needle. a) A series of optical microscope images of four-site implantation of a single flexible probe using a glass needle into the hippocampal region of the mouse brain (i to iv). The black arrows indicate the locations of a single flexible probe that has been implanted. The blue arrows in (iv) indicate the regions of the mesh probe on the brain surface connecting implantation sites I-II, II-III and III-IV. The stereotaxic coordinates of the four implanted sites (I to IV) are as follows (iv): (I): anteroposterior (AP), −2.5 mm; mediolateral (ML), 2 mm; dorsoventral (DV), 2.5 mm, (II): AP, −2 mm; ML, 1.5 mm; DV, 2.5 mm, (III): AP, −2 mm; ML, −1.5 mm; DV, 2.5 mm, and (IV): AP, −2 mm; ML, −2.5 mm; DV, 2.5 mm. Scale bars, 500 μm. b) 3D mapping of the probe-neuron interfaces at four sites (I to IV) in a YFP-H transgenic mouse brain at 2 weeks post-injection. The neurons were genetically modified to express the fluorescent protein YFP. Scale bar, 400 μm. c) 32-channel neural recordings at 2 weeks post-injection. Black, blue, red, and green traces were recorded from site-I, II, III and IV, respectively. d**-**e) Boxplot with individual data points of single-neuron spike amplitude (d) and SNR (e) for each site (black: site-I, blue: site-II, red: site-III, and green: site-IV) from three mice at 2 weeks post-injection. Box plots show mean (blue open squares), median (horizontal lines), quartiles (boxes, 25-75%), and ranges (blue whiskers, 1-99%).

Next, stable electrophysiological performance was verified by recording extracellular action potential spikes in the left and right hippocampal regions of a mouse brain at 2 weeks post-injection (Figure 2c; Figures S6 and S7). In each mouse (N = 3), four-site implantations were performed based on the stereotaxic coordinates with a single probe. Representative 32-channel data recorded from the implanted single probe in the left and right hippocampal regions (I; black, II; blue, III; red, and IV; green in Figure 2c) highlight several key points. First, the connection yield in in-vivo four-site implantations (ca. 88%, N = 3) is statistically similar to that in in-vitro tests (ca. 89%, N = 3; Figure S3) and shows that 7 or 8 electrodes at each of the four sites are capable of recording firing neurons. We note that the lower than expected number of channels recording spiking neurons at site-IV in probes 1 and 3 (Figure S6) reflects incomplete implantation of site-IV into the hippocampal region and not failed electrodes in these probes. Overall, 84 out of 96 channels showed stable recording of individual neurons traces over 4 weeks (Figures S6 and S7). These yield results show that the four-site implantations were achieved without causing damage to the probe.

Second, neural signals were detected at all implantation points (I-IV), and two or three neurons on average were recorded in each channel out of a total of 172 single-unit firing activities at 2 weeks post-injection (Figures S6 and S7). Significantly, all recorded neuron spike waveforms remained consistent over the course of four weeks (Figure S7). These stable waveform results show that the sensing electrodes continued to detect the same neurons at each of the four implantation sites, and consequently, that the probes did not move with respect to the brain tissue at these sites.^13,23^ Third, the average spike amplitudes from four different injection sites (I to IV, N = 3) were ∼49 (I), 44 (II), 47 (III), and 44 μV (IV) (Figure 2d), and signal-to-noise ratios (SNRs) were ∼10 (I), 9 (II), 10 (III), and 11 (IV) at 2 weeks post-injection (Figure 2e). The average spike amplitude was higher in site-I than in site-IV (Figure 2d), although the SNRs were similar due to the higher noise level in site-I (0.32 μV) versus site-IV (0.26 μV). More generally, there is no systematic variation in spike amplitude or SNR for the different sites as shown by comparison of data from three independent mice recorded at 2 and 4-weeks post implantation (Figures S8 and S9). In addition, the total average spike amplitudes of 84 channels were ∼47 μV at 2 weeks and ∼50 μV at 4 weeks post-injection (Figures S8 and S10) with average SNRs of ∼10 at both time points (Figures S9 and S10). The average amplitudes and SNRs show no statistically difference between 2 and 4 weeks post-injection. Thus, the stability of single-neuron recording is confirmed in the four-site implantations of a single probe.

Although in-vivo tests of multiple implantations in four different regions of the brain using a small glass needle were successful, this injection method has two inherent limitations. First, inserting a needle into the brain causes acute damage to the local tissue, and it takes time for the tissues to recover after implantation. Second, the whole probe structure, including the I/O pads that electrically connect individual recording channels in the probe to an external interface, is loaded into a glass needle with a relatively small diameter,^29^ and thus there are constraints on determining the volume and size of the probe. In particular, the glass needle-based injection^22-29^ imposed a limited dimension of the device, making it difficult to further increase the number of electrodes. To overcome these limitations, we propose a new stitching method of flexible electronics based on an ultrathin metal needle akin to a needle and thread used in sewing (Figure 3a). This method has several advantages compared to injection using the glass capillary needle. First, the thickness of the metal needle is ca. 20-35 times smaller than the capillary needle outer diameter, which can reduce substantially acute tissue damage similar to ultrathin Si-probes.^10,17^ Second, this eliminates contraints on the volume and size of the stitching implanted multi-site mesh probes compared with capillary needles.

**Figure 3.**
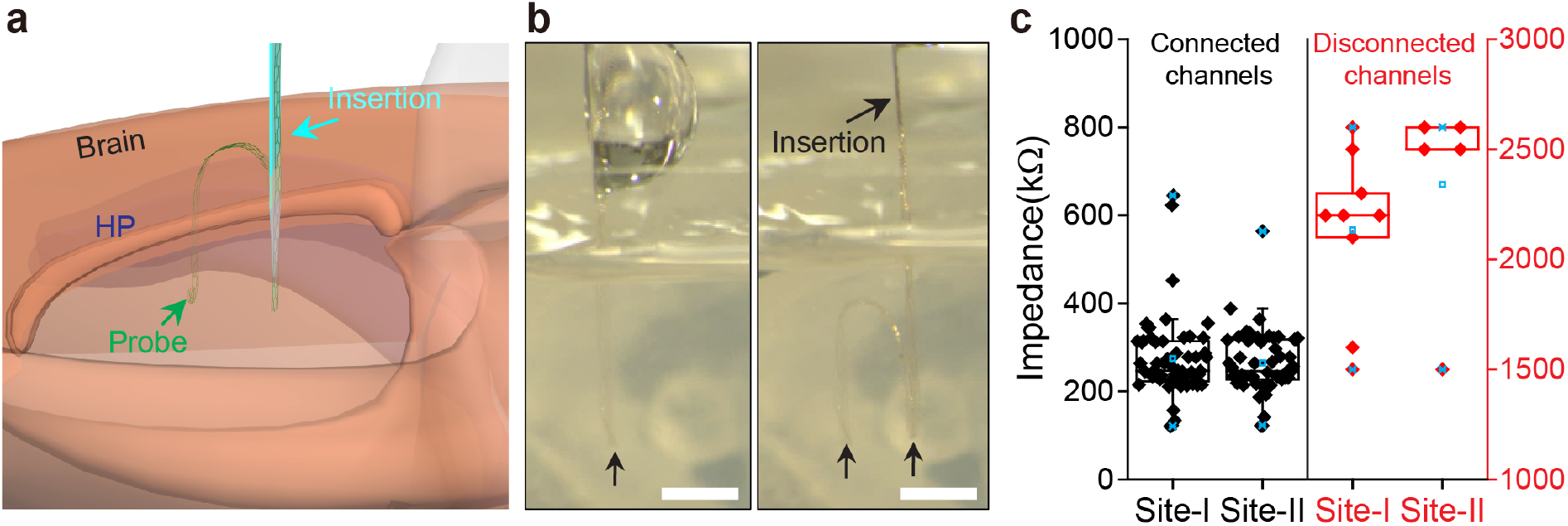
Stitching a flexible probe using a metal insertion needle. a) Schematic of a single flexible probe (green) stitched into the mouse brain via thin metal insertion needle (cyan). b) Optical microscope images (left and right) showing two-site implantation of a single flexible probe into 0.5% hydrogel using metal insertion. Black arrows indicate the implanted sites-I and II in hydrogel. Scale bars, 2 mm. c) In-vitro electrode impedance values at 1 kHz measured from the 32-channel probes (N = 4) implanted in 0.5% hydrogel at two sites (site-I and site-II). Channels that were connected (left, black) and disconnected (right, red) were compared. Box plots show mean (blue open squares), median (horizontal lines), quartiles (boxes, 25-75%), and ranges (blue whiskers, 1-99%).

The implantation method based on metal insertion needle is compatible with standard stereotaxic surgery and does not require specialized equipment. The metal needle,^32^ with an opening angle of ca. 34°, a width of 200 μm and a thickness of 20 μm, was attached to a glass capillary needle with dental cement (Methods and Figure S11). The length of the metal insertion needle varied depending on the target depth of the brain. To minimize damage to the probe during implantation using metal insertion, we designed a new probe structure using finite-element-method (FEM) simulation based on the calculation of von Mises stress distribution^33^ in the probe (Figure S12). The probe with the hexagonal-shaped unit cell versus parallelogram-shaped unit cell of the standard mesh probe was selected to propagate the load stress more evenly along the adjacent probe segments and share the stress with the adjacent unit cells (one length of the hexagon is 47.5 μm) (Figures S12c and S12d).

To examine potential damage to the hexagonal array probe after stitching implantation using the metal insertion needle, we carried out an in-vitro test of implantation in a brain mimicking hydrogel (Figure 3b and Figure S13). Initially, the probe was loaded into a glass capillary needle attached to the top of the metal inserstion needle. There are no restrictions on the size of glass capillary needle because the glass needle is not inserted into the brain in this new method. Next, a small portion of the probe was ejected from the glass capillary needle and adhered to the tip of the metal insertion needle. After positioning and pressing the insertion device over the target location, the probe was inserted into the hydrogel to a desired depth with the metal insertion needle, while the glass capillary needle remained above the surface. Subsequently, the metal insertion needle was retracted, leaving the probe extended in the hydrogel without crumpling (Figure 3b, left). These procedures were repeated at the subsequent insertion site (Figure 3b, right and Movie S2), which also showed that the implantation led to an extended probe structure in the hydogel.

To estimate the connection yield of stitching implantation using the metal insertion needle, we measured impedance the at 1 kHz (Figure 3c) of four 32-channel probes implanted into the hydrogel (N = 4). In these in-vitro tests, 106 out of 128 channels were connected to the FFC (ca. 86% yield) using a criteria that impedence values >1 MΩ corresponded to disconnected channels. Our statistical analysis of the recorded data for the connected channels also shows nearly the same values for sites I and II mean impedence +/- 1SD of 274 +/- 93 and 264 +/- 71 kΩ, respectively. Last, we note that the connection yield is comparable to that determined for multi-site implantion using a glass capillary needle (Figure S3, ca. 89%), suggesting that the hexagonal design was not damaged by insertion with the metal needle.

Having shown that the new hexagonal array probe design and injection method are comparable to the standard mesh design and syringe-based injection method, we further asked whether a stable interface with the neuronal and glial networks is formed during stitching implantation using the metal needle. Several critical points are highlighted in the key steps of the implantation procedure into a live mouse brain (Figure S14). First, a relatively large cranial window (∼3 × 3 mm^2^) was opened to allow for two-site implantations of a single hexagonal array probe based on the stereotaxic coordinates of hippocampal field (anteroposterior: −1.70 mm, mediolateral: −1.60 mm, dorsoventral: 1.17 mm). Second, the dura mater covering the brain was carefully removed through a small incision in the window. This step is important because it is difficult to puncture the dura with the 20 μm-thick metal insertion needle. Third, it is straightforward to implant the hexagonal array probe in at least two sites using the metal insertion needle similar to the in-vitro tests using brain-mimicking hydrogel discussed above.

3D images of tissue sections including the hexagonal array probe implanted in transgenic mice (Thy1-YFP-H and GFAP-GFP) were taken to elucidate the interfaces between the probe and the neurons/astrocytes at 2 weeks post-injection (Figures 4a-b and Figures S15-S16). These images highlight several key points of stitiching implantation with the metal insertion needle. First, acute damage and disturbance of the local tissue were significantly reduced during implantation because the cross-sectional area of the metal insertion needle (4,000 μm^2^) was ca. 50 times smaller than that of the glass capillary needle in Figure 2 (196,250 μm^2^). Second, liquid fluid assistance was not required in the metal insertion-based implantation of the hexagonal array probe. The images show that the probes were linearly elongated without crumpling and delivered into the tissue at the desired location and depth (Figure 4a and Figure S14). Third, the images show that probes do not disrupt the brain tissue; that is, they are seamlessly integrated in a manner similar to the standard mesh structure^29^ despite the slightly higher bending stiffness of the hexagonal array probe.

**Figure 4.**
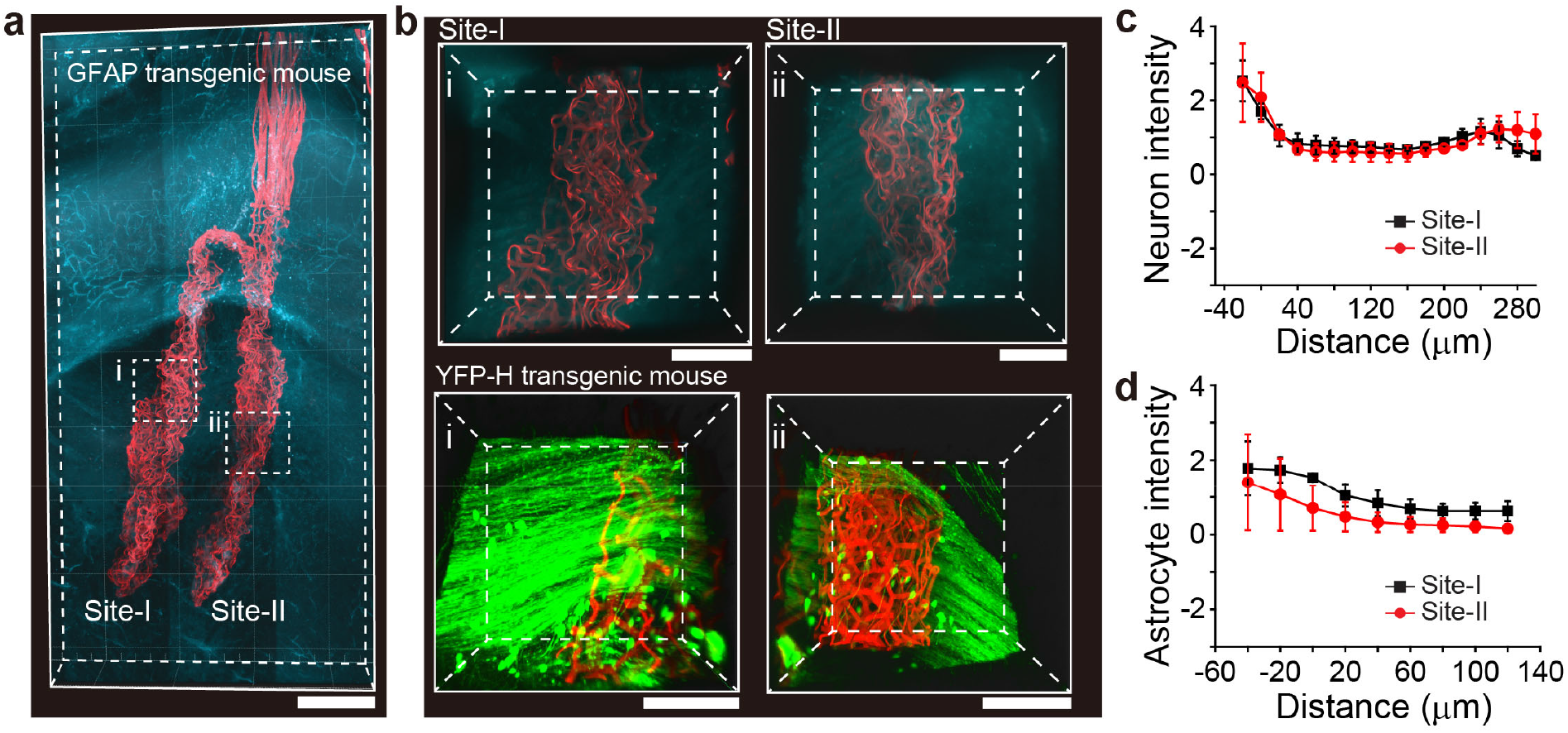
3D mapping and characterization of the probe-neural interface. a) 3D mapping of the probe (red) and astrocytes (cyan) interface in the brain of a GFAP transgenic mouse at 2 weeks post-injection. The astrocytes are genetically modified to express the fluorescent protein. Scale bar, 300 μm. b) High-resolution images of the volumes highlighted by white dashed boxes (i, site-I and ii, site-II) in (a). Scale bars, 100 μm. (Bottom) 3D mapping showing the interface of the probe (red) and neurons (green) in a YFP-H transgenic mouse brain at 2 weeks post-injection. High-resolution images of the volumes highlighted by white dashed boxes (i, site-I and ii, site-II) in Figure S15. Scale bars, 100 μm. c-d) Normalized fluorescence intensity of neurons (c) and astrocytes (d) plotted as a function of the distance from the probe surface site-I (black) and site-II (red) at 2 weeks post-injection. The error bars denote s.e.m.

To quantify the interface of hexagonal array probes with neurons and astrocytes at the two sequentially-implanted sites, higher-resolution images were taken from multiple independent samples at 2 weeks post-implantation with both Thy1-YFP-H (N = 3, total = 6) and GFAP-GFP (N = 2, total = 4) transgenic mice (Figure 4b, Figures S15b and S16). Statistical analyses of the normalized fluorescence intensity of neurons and astrocytes as a function of the distance from the hexagonal array probe surface (Figure 4c and 4d) show a relatively uniform distribution near the probe with no discernible depletion of neurons and only slight enhancement of astrocytes at sites-I or II. Comparison of the data as a function of distance from the probe surface at both implantation sites further shows similar trends for both the normalized neuron and astrocyte intensities at the two sites, suggesting that implantation with the metal needle does not adversely affect brain tissue at the subsequent (i.e., site-II) implantation sites.

To assess whether the hexagonal array probe can detect the electrical signals of neurons in each of the two implantation regions, recording data were obtained from 32-channel probes in live mice brains at 2 and 4 weeks post-implantation (N = 3, Figure 5a and Figure S17). Several key features are revealed by data from the hexagonal array probes in the 1st and 2nd regions (Figures 5a and b; Figure S17). First, analysis of the data recorded at 2 weeks post-implantation in the three mice detected extracellular action potential firing while also exhibiting a high connection yield at both implantation sites each of the implanted probes: site-I, 93.7; site-II, 89.5% with a total yield of 88 out of 96 channels or ca. 91.6%. Furthermore, at 4 weeks post-implantation consistent yields were observed in the two regions for the three mice (Figure S17).

**Figure 5.**
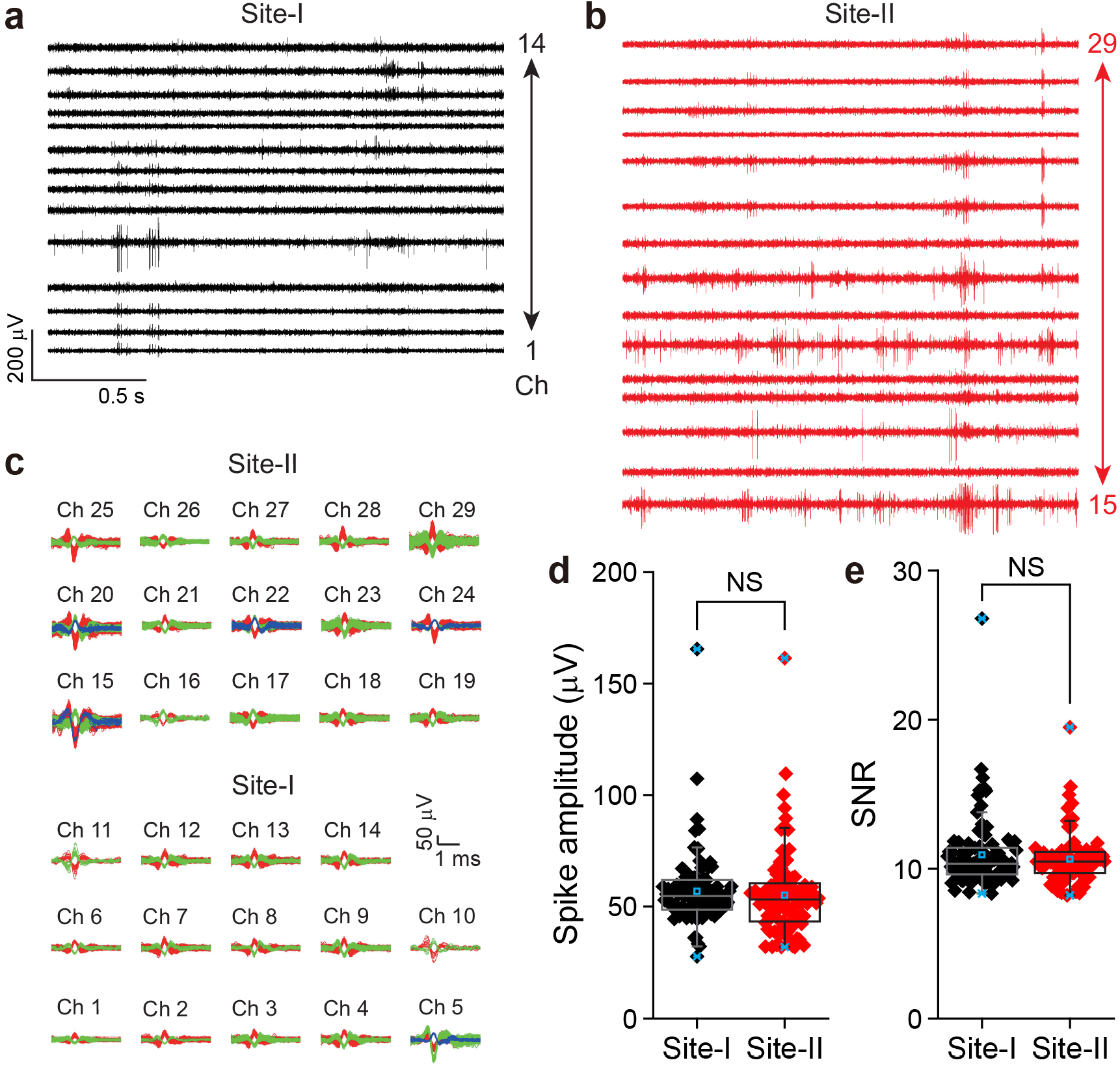
In-vivo recording with a probe implanted in the brain by metal insertion. a) and b) Band-pass (250-6000 Hz) filtered extracellular spike recordings from the site-I (a) and site-II (b) at 2 weeks post-injection. c) Overlay of sorted and clustered spikes from the site-I (Ch 1 to Ch 14, bottom) and the site-II (Ch 15 to Ch 29, top) regions of recording traces in (a) and (b), respectively. Red, green, and blue colors are used to denote spikes assigned to different neurons. d-e) Boxplot of individual data points of single neuron spike amplitude (d) and SNR (e) for each site (black: site-I and red: site-II) from three mice at 2 weeks post-injection. Box plots show mean (blue open squares), median (horizontal lines), quartiles (boxes, 25-75%), and ranges (blue whiskers, 1-99%). NS: not significant (P > 0.05).

Second, waveform analyses of the recorded data (Figure 5c; Figure S18) show that two or three neurons were recorded on average by each channel at 2 weeks post-implantation. 29 and 34 single unit neurons were recorded site-I and site-II, respectively (Figure 5c). In addition, the total number of recorded single units are nearly unchanged over the 4 weeks of recording: 189 single-unit spikes at 2 weeks vs. 188 single-unit spikes at 4 weeks (Figure S18). This confirms the stability of single-neuron activities recorded by the hexagonal array probe, and shows that the probes did not move with respect to the brain tissue at sites-I and II. Moreover, the average amplitudes of the sorted single-unit spikes at 2 weeks were ca. 56 μV (45 channels) and 55 μV (43 channels) at site-I and site-II, respectively, and remained similar at 4 weeks (Figure 5d and Figure S19). The average SNRs were also nearly the same (ca. 10) for single units recorded at both sites at 2 weeks post-injection (Figure 5e). Furthermore, the SNRs remained nearly the same even at 4 weeks post-implantation (Figure S20). Hence, the quantification of spike amplitudes and SNRs revealed no statistically significant difference between at sites-I and-II, and are consistent with a stable probe/brain interface for hexagonal array probes implanted at least two sites via the metal insertion needle methodology.

## 3. Conclusion

In conclusion, we successfully demonstrated multi-site stitching implantation of ultra-flexible and biocompatible single probes using glass capillary and ultrathin metal insertion needles, and subsequent monitoring of neural spiking from 2 or 4 distinct brain sites using a single low-mass interface. To facilitate four-site implantations using glass capillary needle, the design and implantation process of the flexible probe were optimized through in-vitro tests using hydrogel, after which the connection yields were quantitatively evaluated (ca. 89%). The statistical analysis of in-vivo histology images and recording of electrophysiology data at each implantation site showed seamless interface and chronic stability over one month. In addition, we introduced a new implantation method using a ultrathin metal insertion needles that are analogous in many ways to sewing with a needle and thread. A hexagonal-shaped array probe was designed with reference to FEM simulations. In-vitro implantation tests into hydrogel were also performed to validate the injection process and evaluate connection yields (ca. 86%). Notably, three-dimensional immunohistochemical staining and imaging at each implantation site highlighted the seamless and stable hexagonal-shaped array probe. Electrophysiology experiments were carried out and spike sorting data were analyzed, and demonstrated the ability to track in parallel single neuron activities across each injection site. These data and analyses of the temporal dynamics of average spike amplitudes and SNRs further showed chronic stability for at least a 1-month time scale. While the implantation method using a metal insertion needle requires the removal of the dura in the region of injection site, it has been shown to minimize acute damage to the local tissue during implantation, overcoming limitations in the size and volume of the probe and allowing for the possibility of increasing the number of recording channels. In the future, it will be interesting to explore implantations where each site consists of a small craniotomy, and the probe is passed over the top of the skull between sites to minimize the area of dura removed during the surgery.

Overall, multi-site implantation of flexible mesh electronics into the brain via stitching using a smaller delivery shuttle opens up a new paradigm for flexible probe design and multi-site implantation in the brain and possibly other tissues. We believe that this approach could provide critical information in fundamental neuroscience research,^34^ such as full understanding of learning^35^ and memory storage^36^ by chronically monitoring thousands of single-unit level neural signals from multiple targeted sites of the brain, be adapted to wireless interfaces that eliminate constraint of cables to the recording electronics,^20,21^ be applicable for implantation of ultraflexible electronics probes for therapeutic purposes, and could be applied to other classes of flexible electronics.

## 4. Experimental Section/Methods

### Design and fabrication of flexible probes

Two different probes were examined depending on whether a glass needle or a metal insertion needle was used (standard mesh vs. hexagonal array probe). Key parameters of both device structures are as follows: (i) total mesh widths are W_n_ = 1 mm for standard mesh and W_m_ = 1.2 mm for hexagonal array probe; (ii) all SU-8 ribbon widths are 10 μm for both designs; (iii) the angles between longitudinal and transverse SU-8 ribbons are α_1_ = 45° for standard mesh and α_2_ = 120° for hexagonal array probe; (iv) longitudinal spacing (pitch between transverse ribbons) is L_n1_ = 333 μm and transverse spacing (pitch between longitudinal ribbons) is L_n2_ = 62.5 μm for standard mesh, and the one-side length of the hexagonal unit cell is L_m1_ = 47.5 μm for hexagonal array probe; (v) all metal interconnect line widths are 1.5 μm and the total number of recording channels is 32 for both designs; (vi) longitudinal ribbons have a three-layer sandwich structure (SU-8 polymer/metal/SU-8 polymer) and transverse ribbons have a two-layer structure (SU-8 polymer/SU-8 polymer) for both designs. The fabrication steps of both mesh structures were similar to those of the previous reports.

### Preparation of hydrogel

0.5 g agarose (SeaPlaque® Lonza Group Ltd) was mixed with 10 mL 10× TAE (Tris-Acetate EDTA) and 90 mL DI water in a glass beaker. The beaker was covered with a piece of aluminum foil (Reynolds Wrap® Reynolds Consumer Products) to prevent evaporation, and heated at boiling on a hot plate until the solution was clear: the final mass concentration was ca. 0.5%. The solution was allowed to naturally cool to room temperature after pouring the solution into a glass container for in-vitro implantation tests (Figures 1c and 3b; Movies S1 and S2).

### Vertebrate animal subjects

Adult (20-30 g) female CD-1 mice (Charles River Laboratories, Wilmington, MA) were used for electrophysiology recording of stitching mesh electronics as vertebrate animal subjects in this study. B6.Cg-Tg(Thy1-YFP)HJrs/J and FVB/N-Tg(GFAPGFP)14Mes/J transgenic mice (The Jackson Laboratory, 4-6 weeks old) were used for histology analyisis and study. All procedures performed on the vertebrate animal subjects were approved by the Animal Care and Use Committee of Harvard University. The animal care and use programs at Harvard University meet the requirements of the Federal Law (89-544 and 91-579) and NIH regulations and are also accredited by the American Association for Accreditation of Laboratory Animal Care (AAALAC). Animals were group-housed on a 12 h: 12 h light: dark cycle in the Harvard University’s Biology Research Infrastructure (BRI) and fed with food and water ad libitum as appropriate.

#### Histological sectioning and imaging

##### 1. Sample preparations

The mesh probes and hexagonal array probes were labeled with Lissamine Rhodamine B ethylenediamine prior to surgical injection. 10 μg Lissamine Rhodamine B ethylenediamine were added to 50 mL SU8-2000.5 and vigorously stirred for 3 days in the dark, followed by centrifuging at 6000 rpm to remove any Lissamine Rhodamine B ethylenediamine residue. The SU8-2000.5 was then used for the fabrication of labeled mesh probes.

Mice with implanted labeled mesh probes were anaesthetized with ketamine/dexdomitor in saline, perfused with 40 mL ice-cold Phosphate-buffered saline (1×, HyClone) and subsequently with 4% paraformaldehyde (Electron Microscopy Sciences) delivered at 2 ml/min, and decapitated. After removing the scalp skin, the skull was thinned using a high-speed rotary grinder and gently removed using micro-scissors and tweezers. Brains were then immersed in 4% paraformaldehyde for 24 h, and transferred to 1× PBS for another 24 h to remove unreacted paraformaldehyde. After embedding in 4% agarose hydrogel (SeaPlaque agarose, Lonza) in 1× PBS, the brains were cut into 2 cm (length) × 2 cm (width) × 1 cm (height) blocks. The meshes in the blocks were observed by micro-CT X-ray scanning (HMX ST 225, Nikon Metrology). CT Pro 3D software (Nikon Metrology) was used to reconstruct the observed 3D images, and VGStudio MAX 3.0 software (Volume Graphics) was used to render and analyze the 3D reconstructed images.

The mouse brain was sectioned into 250-800 μm slices using a vibratome (VT1000S vibrating blade microtome, Leica). The slices were incubated in the refractive index matching solution composed of glycerol(Sigma-Aldrich)/PBS (80%/20%) for 24 hours, and transferred to a 50-mm-diameter Petri dishes and glued using Devcon epoxy (ITW Polymers Adhesives) at their edges. After epoxy curing, 30 mL of glycerol (Sigma-Aldrich)/PBS (80%/20%) was added to the petri dish to ensure that the brain slices were fully immersed.

##### 2. Fluorescence imaging

Fluorescence imaging was conducted using a confocal/multiphoton microscope (Carl Zeiss Microscopy LSM 880) with a 20× objective (numerical aperture 1.0, free working distance 5.6 mm). Images were acquired under either confocal or 2-photon mode. For confocal imaging, lasers with wavelengths of 405 nm, 488 nm and 561 nm were used to excite endogenously expressed YFP and GFP in transgenic mouse brain slices and Lissamine Rhodamine B ethylenediamine in mesh probes. For two-photon imaging, near-infrared lasers with wavelengths of 950 nm, 920 nm and 840 nm were used to excite YFP, GFP, and Lissamine Rhodamine B ethylenediamine, respectively. The bandpass filters with wavelengths of 575-610 nm and 500-550 nm were used for the fluorecense measurements of Lissamine Rhodamine B ethylenediamine and YFP, respectively. Zen Blue (Zeiss) and Imaris (Oxford Instruments) softwares were used to stitch and visualize the images, respectively.

### Numerical simulations of von Mises stress distribution

We performed finite-element method (FEM) simulations to calculate structural deformations and von Mises stress distributions (COMSOL Multiphysics; Figure S12a-c). Three mesh structures with a parallelogram unit cell and two different-sized hexagonal unit cells were examined. Detailed structural parameters were shown in Figure S12. A thin rod with a size of 200 × 20 × 340 μm^3^ was pressed down to the depth of 10 μm at the center of each mesh structure, to apply a local deformation. The built-in contact pressure function in COMSOL was used to simulate the local deformation. The effect of the rod orientation was examined by calculating structural deformations and von Mises stress distributions when the rod was parallel to the horizontal direction and perpendicular to the oblique side of the unit cell. The mass density, Young’s modulus, and Poisson ratio of SU-8 were set to the values of 1190 kg/m^3^, 4.02 GPa, and 0.22, respectively. A fixed boundary condition was imposed at the end of the full-domain mesh structures in the FEM simulation.

#### Key steps of mouse surgery

##### 1. Survival mouse surgery of stitching mesh electronics using glass capillary needle

∼3 × 2 mm^2^ window was carefully drilled in the mouse skull by dental drill (Micromotor with On/Off Pedal 110/220, Grobet USA) for four-site implantations. The 30 mm-length mesh probe was loaded into the needle (ID: 300 μm, OD: 450 μm). Using the field-of-view (FoV) injection method, the first implantation was carried out with an injection rate of 5-10 mL/h by a syringe pump (PHD 2000, Harvard Apparatus). From the second implantation, the surgery was performed without the aid of liquid flow in both hemospheres. Sufficient mesh probe is ejected or pulled from capillary prior following the first or subsequent implantation sites so that excessive stress is not placed on the mesh and brain tissue during implantation at the next site.

##### 2. Survival mouse surgery of stitching mesh electronics using metal insertion

∼3 × 3 mm^2^ of skull was opened by dental drill and the dura was removed by a syringe needle. A metal insertion needle with 200 μm-width, 20 μm-thickness, and ∼34°-opening anlge was attached to a glass needle (ID: 600 μm, OD: 750 μm) by dental cement. The whole hexagonal array probe was loaded into the needle. Then, ∼1 mm-long mesh tip was positioned underneath the metal insertion needle so that the hexagonal array probe was delivered into the brain by pushing the metal insertion needle. The metal insertion needle was lifted up by motorized stereotaxic frame and moved to other target area. Lastly, the I/O pads of the mesh were aligned to FFC conductors for electrical connection by direct contact method.

### In-vivo recording in the mouse brain

Mice with multi-site implantations were restrained in a Tailveiner restrainer (Braintree Scientific LLC). Electrophysiological recording was performed using Intan evaluation system (Intan Technologies LLC) with a 20-kHz sampling rate and a 60-Hz notch filter.

### Analysis of electrophysiological recording data

The extracellular electrophysiological recording data were analyzed offline. Briefly, raw recording data were filtered using noncausal Butterworth band-pass filters (“filtfilt” function in Matlab) in the 250- to 6000-Hz frequency range to extract single-unit spikes. Single-unit spike sorting (Figure 5c; Figures S7 and S18) was carried out by amplitude thresholding of the filtered traces, where the threshold was automatically selected based on the median of the background noise according to the improved noise estimation method. The average spike amplitude for analyzed channels (Figures 2d and 5d; Figures S8 and S19) was defined as the average peak-to-peak amplitude of all recorded spikes in those channels. In addition, to obtain the noise level in each channel, the median of the absolute values of the recorded traces was divided by 0.6745, which provides reasonable approximation of 1 standard deviation of the noise distribution excluding firing events. The median of the recorded traces was used instead of the standard deviation, to minimize the influence of relatively high amplitude firing events. The signal-to-noise ratio (SNR) was then calculated in each channel (Figures 2e and 5e; Figures S9 and S20) by dividing the average spike amplitude by the corresponding noise level.

## Acknowledgements

C.M.L. acknowledges partial support of this work by the Air Force Office of Scientific Research (FA9550-14-1-0136). H.-G.P. acknowledges support from a National Research Foundation of Korea (NRF) grant funded by the Korean government (MSIT) (2021R1A2C3006781).

## Author contributions

J.M.L., H.-G.P. and C.M.L designed the experiments. J.M.L., and D.L. performed the experiments. J.M.L. and H.-R.K. performed the simulations. J.M.L., D.L, Y.-W.P, H.-G.P. and C.M.L analyzed the data. J.M.L., D.L., H.-G.P. and C.M.L wrote the paper. All authors discussed the results, revised, or commented on the manuscript. H.-G.P. and C.M.L supervised the project.

