## Supplemental Information for "Stitching flexible electronics into the brain"

**This file includes:**

Supplementary Figures 1 to 20

Supplementary Movie Legends 1 to 2

### Supplementary Figures

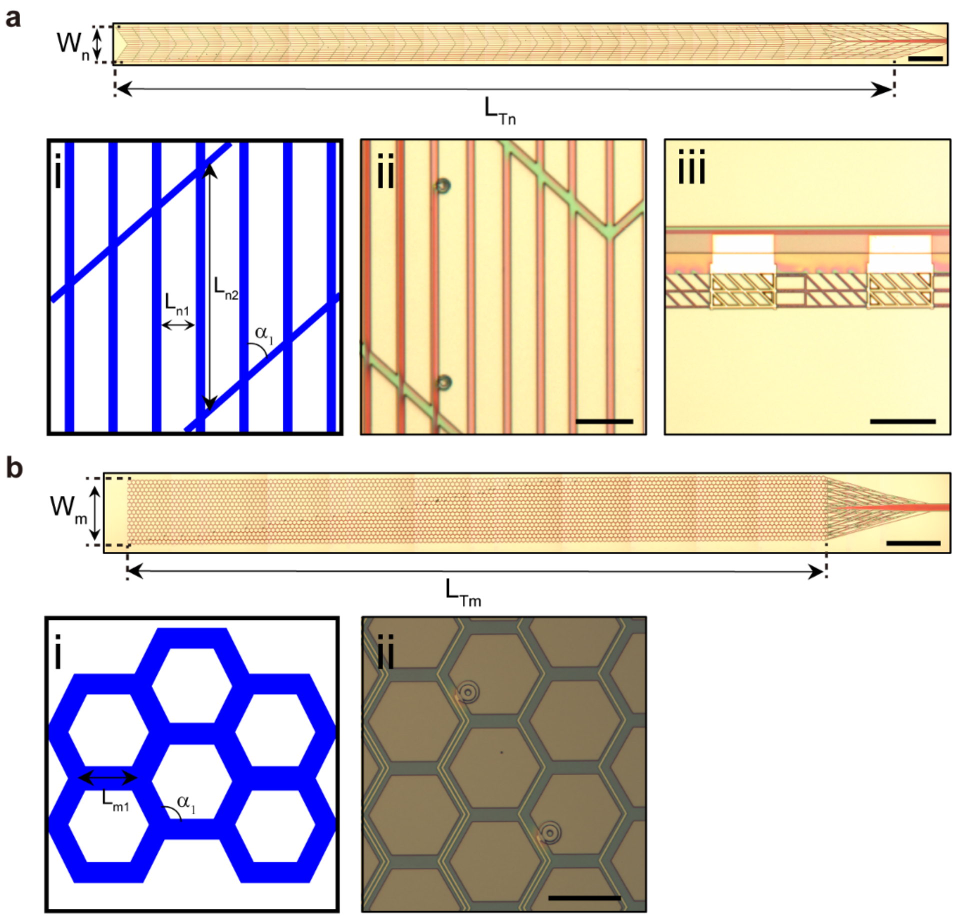


**Figure S1. Overall design of stitching flexible electronics for multi-site implantations using glass capillary and metal insertion needles. a**, Schematic of a 32-channel flexible probe for four-site implantations using glass capillary needle. The total width of probe, $\mathbf{W_n}$, and the total length of probe, $\mathbf{L_{Tn}}$, are 1 mm and 30 mm, respectively. Scale bar, 1 mm. **b**, (i) Schematic of a portion of the probe for four-site implantations. The widths of transverse and longitudinal elements are 10 μm; angle between transverse and longitudinal elements, $\boldsymbol{\alpha_1}$ = 45°; pitch in longitudinal direction, $\mathbf{L_{n1}}$ = 330 μm; and pitch in transverse direction, $\mathbf{L_{n2}}$ = 62.5 μm. (ii)-(iii) Optical microscope images of the mesh and I/O pad parts. All Pt-coated recording electrodes terminate at individual I/O pads. Scale bars, 100 μm (ii) and 200 μm (iii). **c**, Schematic of a 32-channel probe for multi-site implantations using an micron thickness metal insertion needle. The total width of mesh, $\mathbf{W_m}$, and the total length of mesh, $\mathbf{L_{Tm}}$, are 1.2 mm and 15 mm, respectively. Scale bar, 1 mm. **d**, (i) Schematic of a portion of the mesh for two-site implantations. All element widths are 10 μm; angle between two elements, $\boldsymbol{\alpha_2}$ = 120°; one-side length of the hexagonal unit cell, $\mathbf{L_{m1}}$ = 47.5 μm. (ii) Optical microscope images of the recording electrode portion of the probe. All Pt-coated recording electrodes terminate at individual I/O pads as shown in (a, iii). Scale bar, 50 μm.

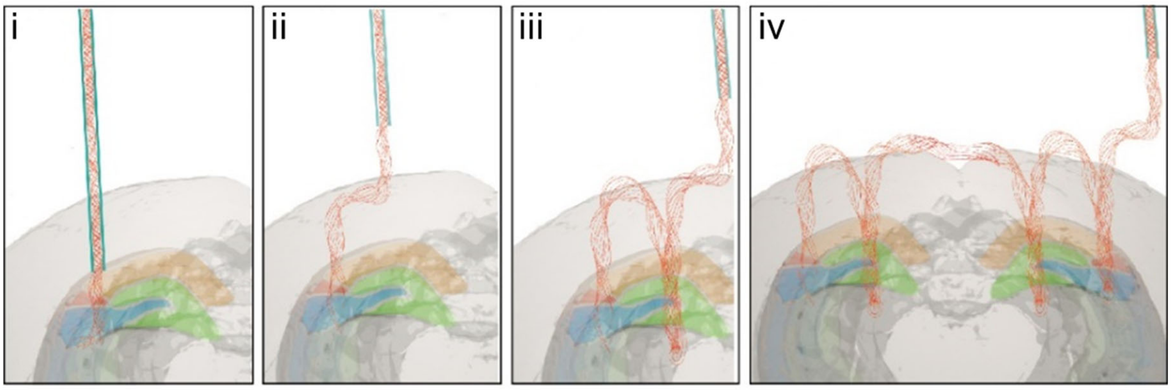


**Figure S2. Schematics illustrating key steps of four-site implantations of a single flexible probe.** A series of schematics (i to iv) of the probe being implanted into four different regions of mouse brain. (i) The probe (red) is loaded in a glass capillary needle with sterile saline and stereotaxically implanted into the left hippocampus. (ii) The glass capillary needle is withdrawn and moved to the next target position using the motorized stereotaxic frame. (iii and iv) The first and second processes are repeated in the next insertion sites including target areas in the right hippocampus.

In the case of implantation using a glass capillary needle (ID 0.30, OD 0.45, and length 100 mm capillaries were used in our studies), the minimal and maximum distances between regions of implantation are ~0.3 and ~60 mm, respectively. Given that the capillary needle has a wall thickness of 0.15 mm, the theoretical next-nearest injection site would be ~0.3 mm away from the first one, although we have not tried to experimentally determine this value. To ensure a smooth loading and unloading process, the length of the probe should not exceed the 100 mm length of the capillary needle. For 2-site implantations with implantation depths of 3-5 mm (i.e., the depth used in the present mouse studies) and a conservative 25 mm left for I/O connection, the maximum site separation would be ~60 mm. However, this maximum separation could be increased using a longer capillary needle. During surgery, we made a ~3 × 2 $mm^2$ craniotomy to access the four different implantation sites (Figure 2a, i-iv). It is also possible to open four smaller craniotomies and pass the probe from the top of the scull to access different areas of the brain, when the areas of interest are far apart or there are potential risks associated with a larger craniotomy.

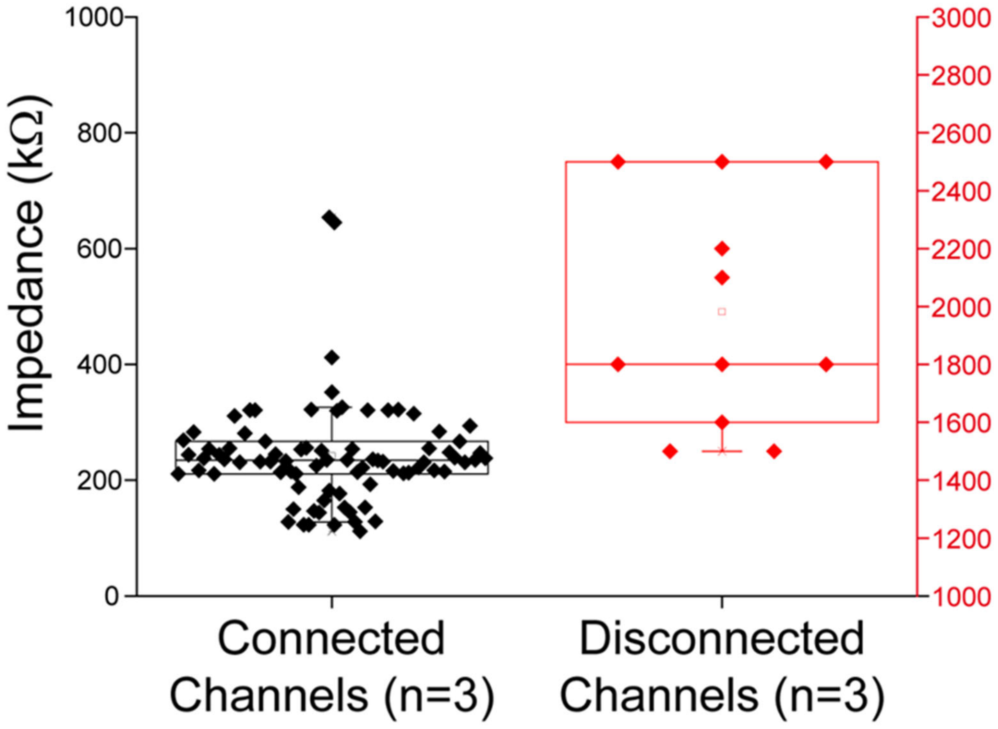


**Figure S3. In-vitro connection yields following four-site implantations. a**, Measured impedance values at 1 kHz for three 32-channel meshes following four-site implantation into brain tissue-like hydrogel.[30,31] Implantation was carried out by sequential injections of the continuous probe through a 300 μm ID glass needle. Summary of the electrode impedance values for the three implanted probes (N = 3). The bar and whisker plots show individual data points of impedances (96 channels). Box plots show mean (open squares), median (horizontal lines), quartiles (boxes, 25-75%), and ranges (whiskers, 1-99%).

Using a criteria of impedance values less than 1 MΩ as the definition of successful interface connection defines connection yields of ~91%, ~88%, and ~88% for the three 32-channel meshes with direct contact I/O interfaces after four-site implantations into hydrogel.

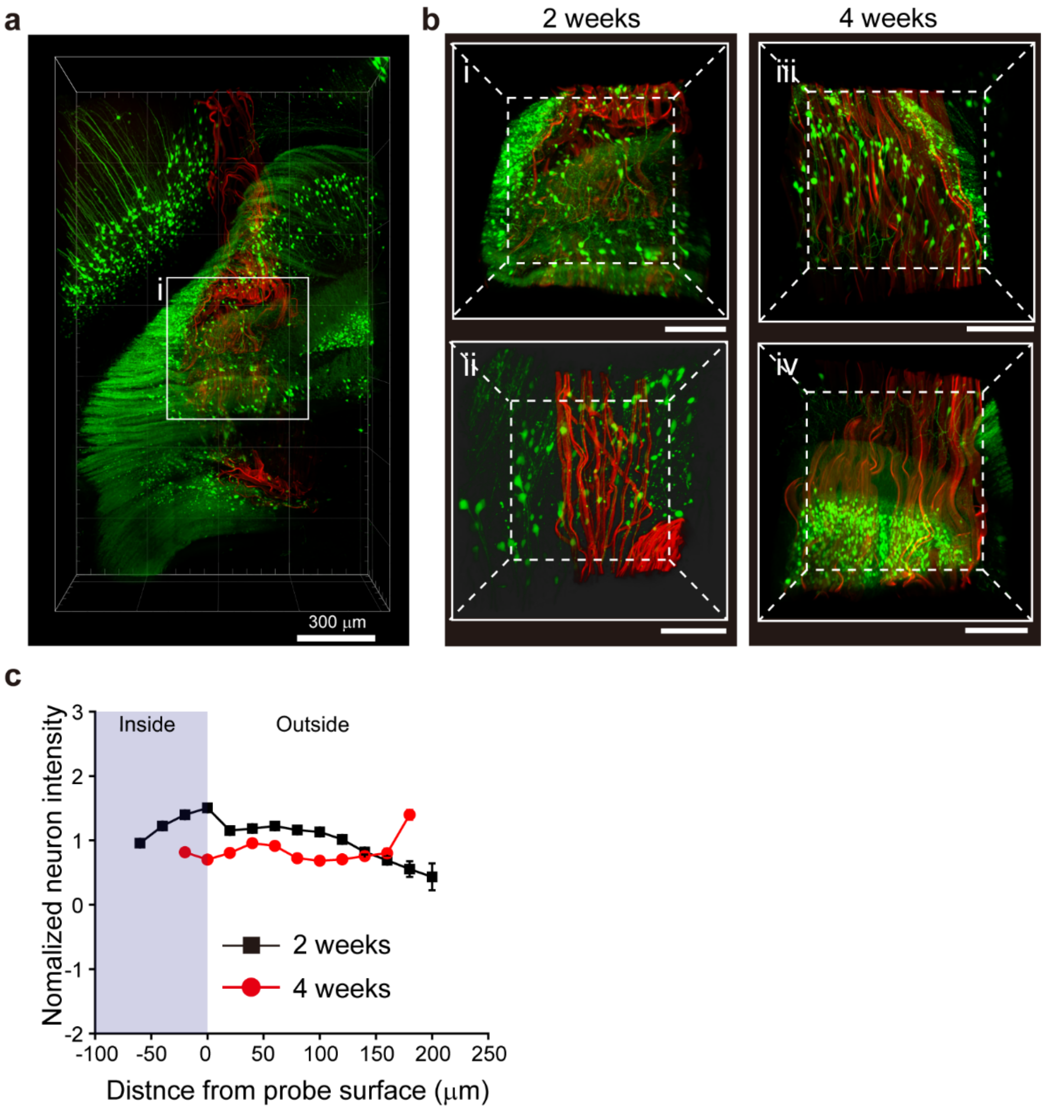


**Figure S4. High-resolution 3D mapping and analysis of probe-neural interface. a**, 3D full probe interfaces between neurons (green) and probe (red) at 2 weeks post-injection. Scale bars, 300 μm. **b**, High-resolution image of the volumes highlighted by white dashed boxes in **a** (i and iii) and high-resolution image from the other YFP-H transgenic mouse brain (ii and iv) at 2 weeks (left column) and 4 weeks (right column) post-injection. Scale bars, 100 μm. **c**, Normalized fluorescence intensity of neurons as a function of distance from the probe boundary at hippocampus at 2 weeks (black) and 4 weeks (red) post-injection. The purple-shaded regions indicate tissue volumes within the interior of the probe. The relative signal was obtained by normalizing the fluorescence intensity by the baseline value defined as the average fluorescence intensity of 3D mapping image.

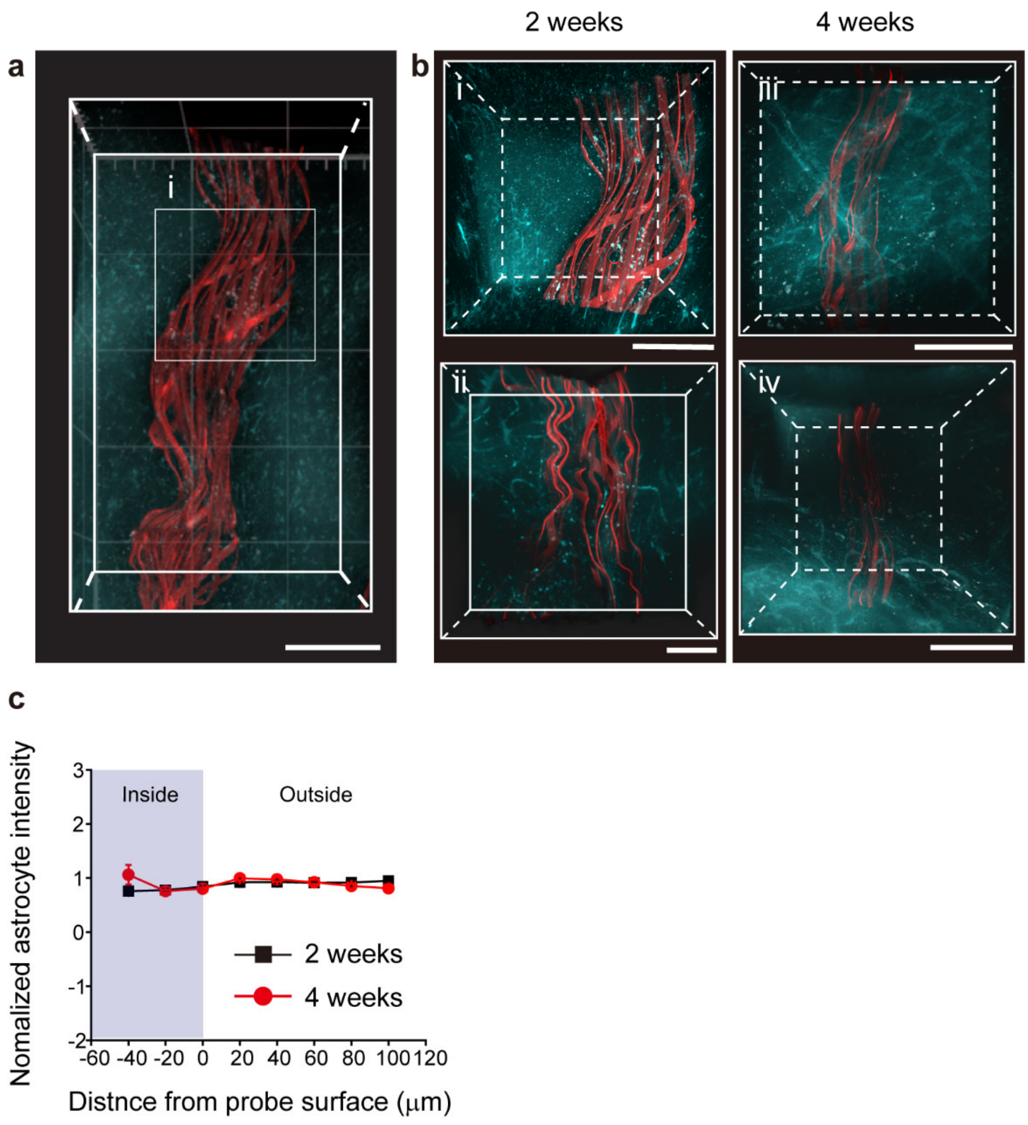


**Figure S5. High-resolution 3D mapping and analysis of probe-astrocyte interface. a**, 3D full probe interfaces between astrocytes (cyan) and probe (red) at 2 weeks post-injection. Scale bars, 300 μm. **b**, High-resolution image of the volumes highlighted by white dashed boxes in **a** (i and iii) and high-resolution image from the other GFAP transgenic mouse brain (ii and iv) at 2 weeks (left column) and 4 weeks (right column) post-injection. Scale bars, 100 μm. **c**, Normalized fluorescence intensity of astrocytes as a function of distance from the probe boundary at hippocampus at 2 weeks (black) and 4 weeks (red) post-injection. The purple-shaded regions indicate tissue volumes within the interior of the probe. The relative signal was obtained by normalizing the fluorescence intensity by the baseline value defined as the average fluorescence intensity of 3D mapping image.

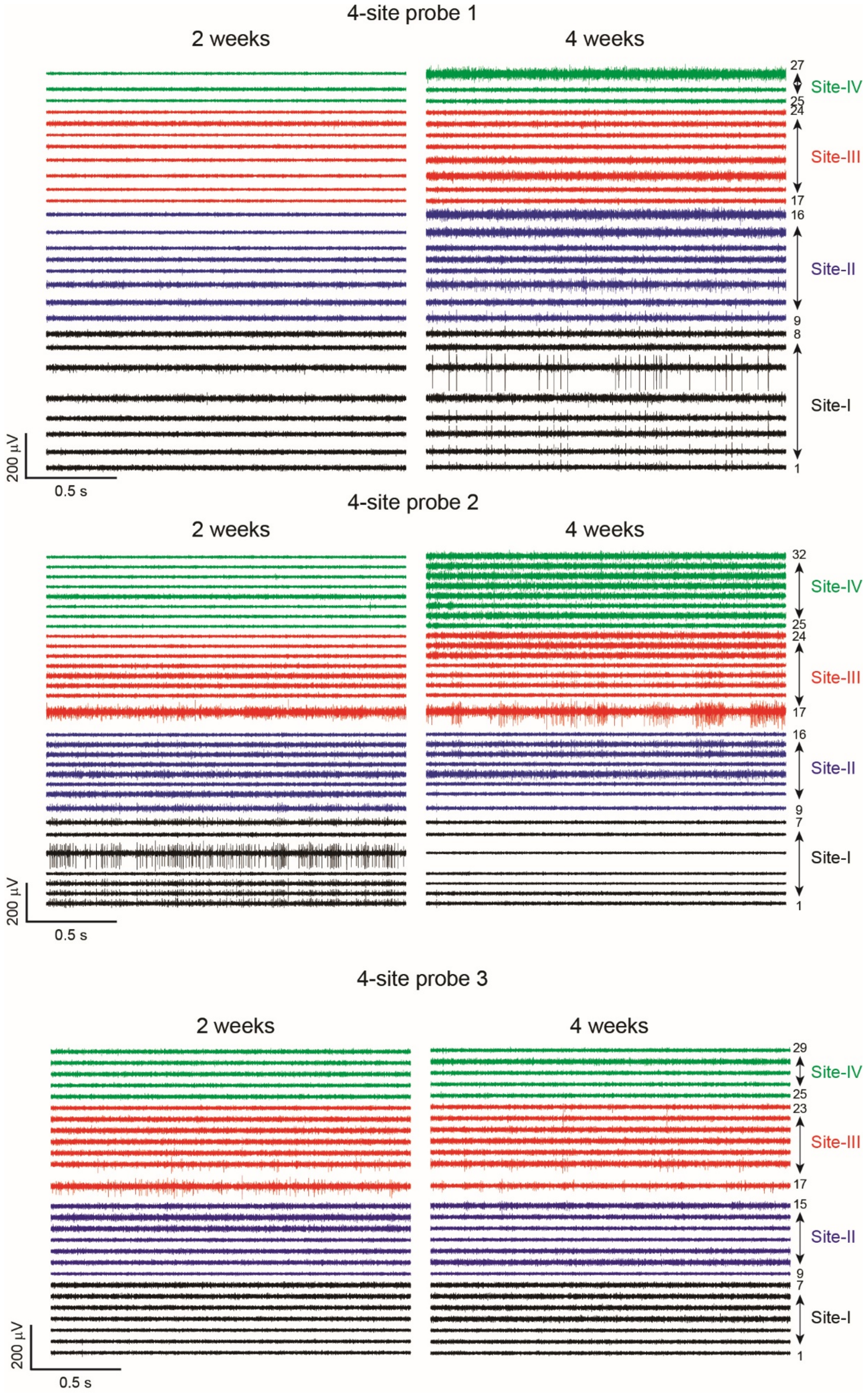


**Figure S6. Extracellular recordings by mesh probe implanted using glass capillary needle.** Band-pass (250-6000 Hz) filtered extracellular single-unit spike traces recorded after four-site implantations in three individual mice (N = 3) at 2 and 4 weeks post-injection. Black, blue, red, and green traces were recorded from site-I, II, III, and IV, respectively. All channels showing neuronal spiking are included in the traces. The probe design (Figure 1b) places 8 addressable electrodes in the regions designed to be implanted at the four sites, and typically, there are 7 or 8 electrodes with impedance values (< 1 MΩ) capable of reliable recording of spiking neurons. The lower number of channels showing single unit spiking at site-IV in probes 1 and 3 reflects incomplete implantation into the hippocampal region of the mouse brain and not failed electrodes.

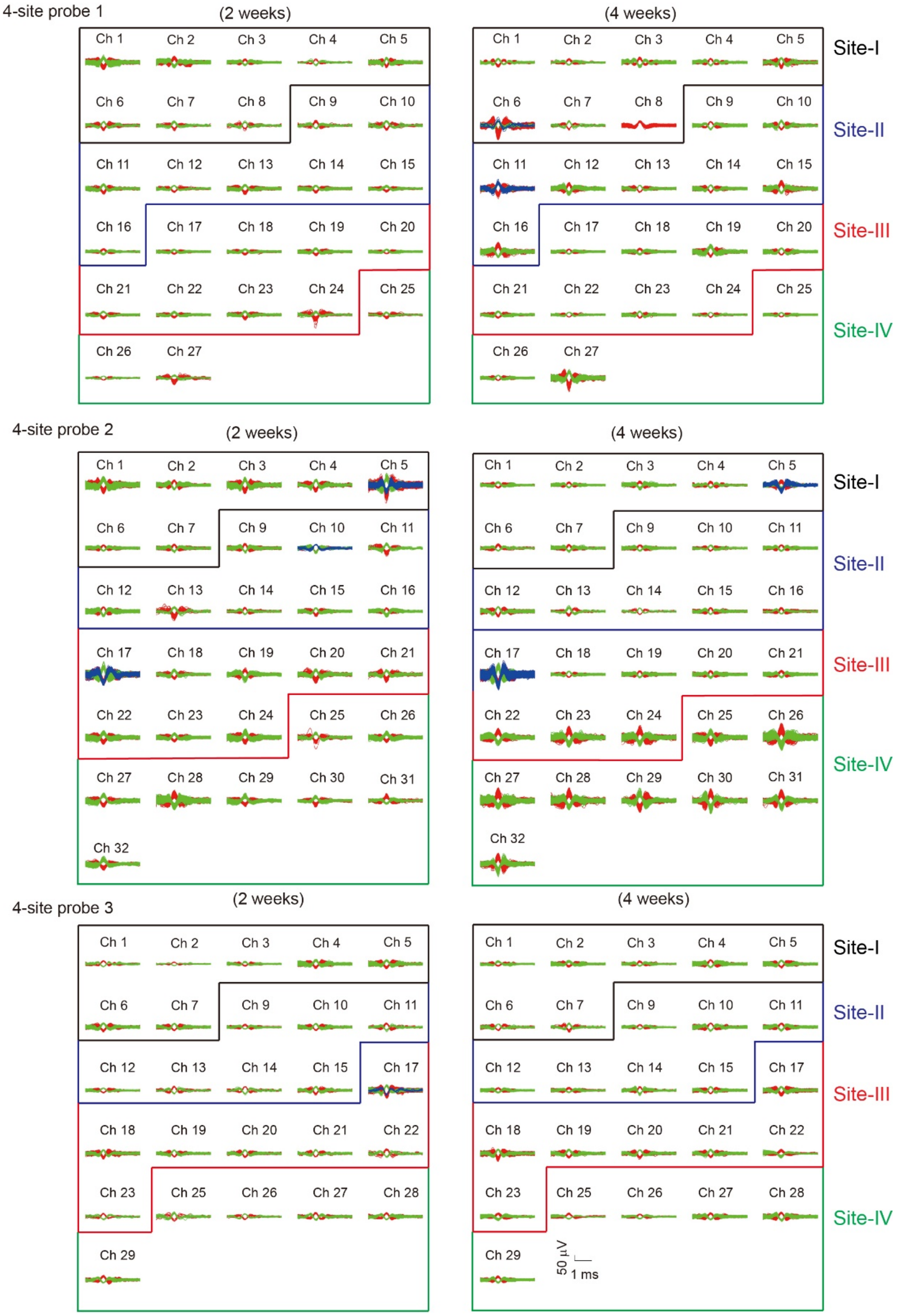


**Figure S7. Overlay of sorted and clustered spikes. a-c**, Spike sorting analysis of the recording data shown in Figure S6 (total 84 channels in three mice at 2 and 4 weeks post-injection). There are 45 (site-I), 47 (site-II), 48 (site-III), and 32 (site-IV) single-unit spikes from 84 channels at

2 weeks; 45 (site-I), 47 (site-II), 49 (site-III), and 32 (site-IV) single-unit spikes from 84 channels at 4 weeks.

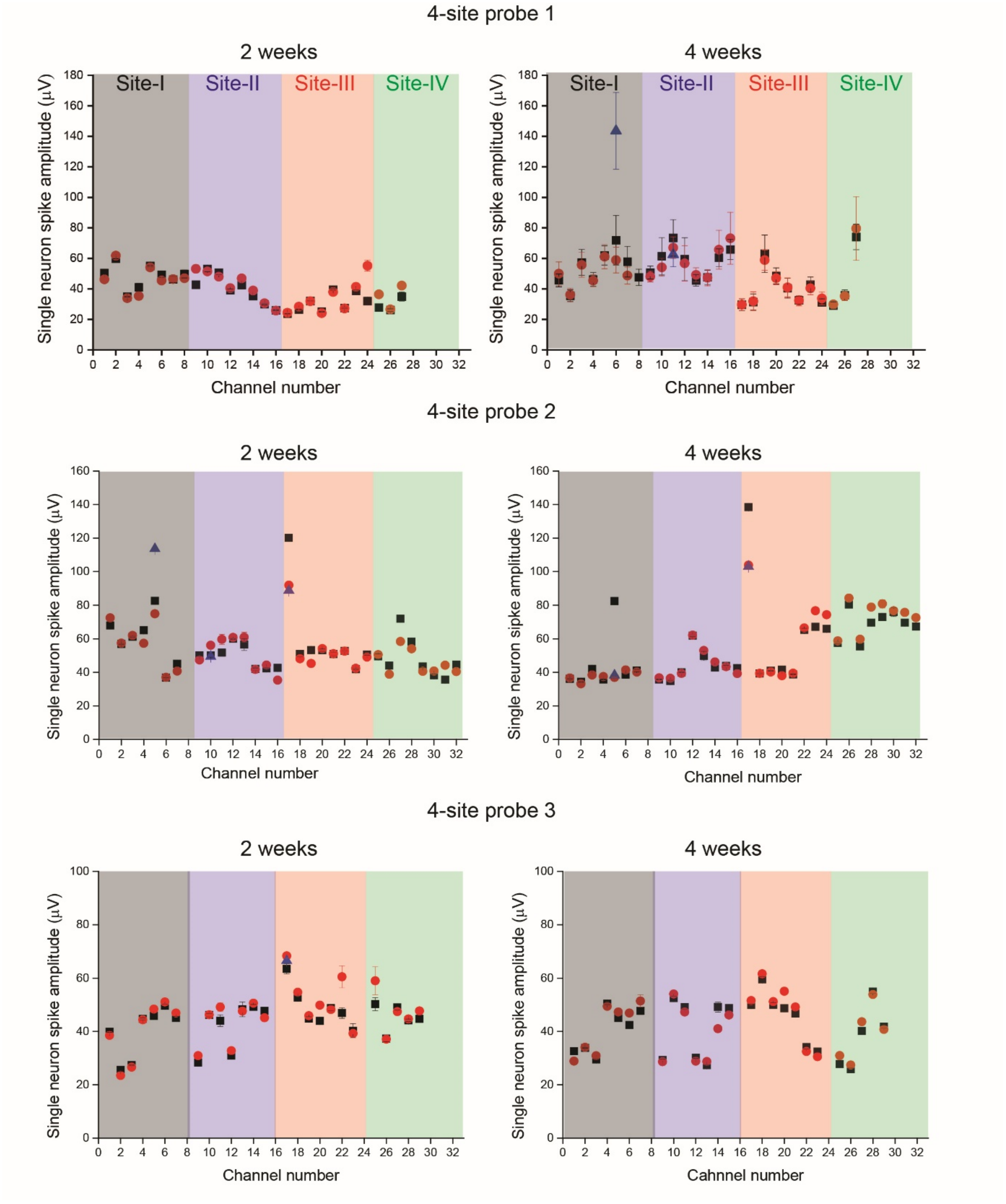


**Figure S8. Spike amplitudes from each implanted site**. **a-c**, Single-unit peak-to-peak spike amplitudes from three mice (N = 3) at 2 and 4 weeks. Data were obtained from Figure S7. The average spike amplitudes from four different injection sites (site-I to IV, N = 3) were ~49 (I), 44 (II), 47 (III), and 44 μV (IV), respectively at 2 weeks post-injection.

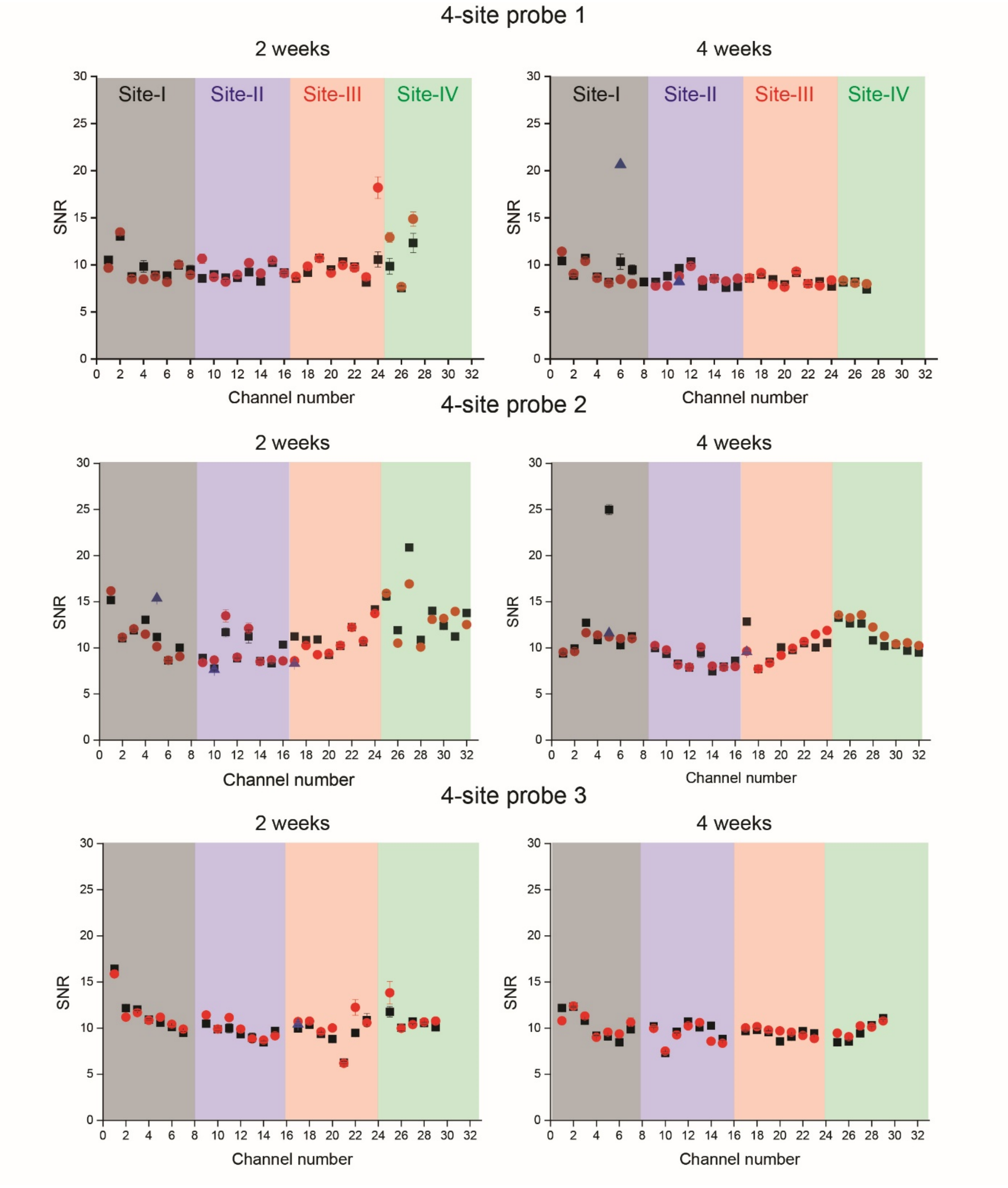


**Figure S9. SNRs from each implanted site. a-c**, Single-unit SNRs from three mice (N = 3) at 2 and 4 weeks. Data were obtained from Figure S7. The average signal-to-noise ratios (SNRs) were ~10 (site-I), 9 (site-II), 10 (site-III), and 11 (site-IV) at 2 weeks post-injection.

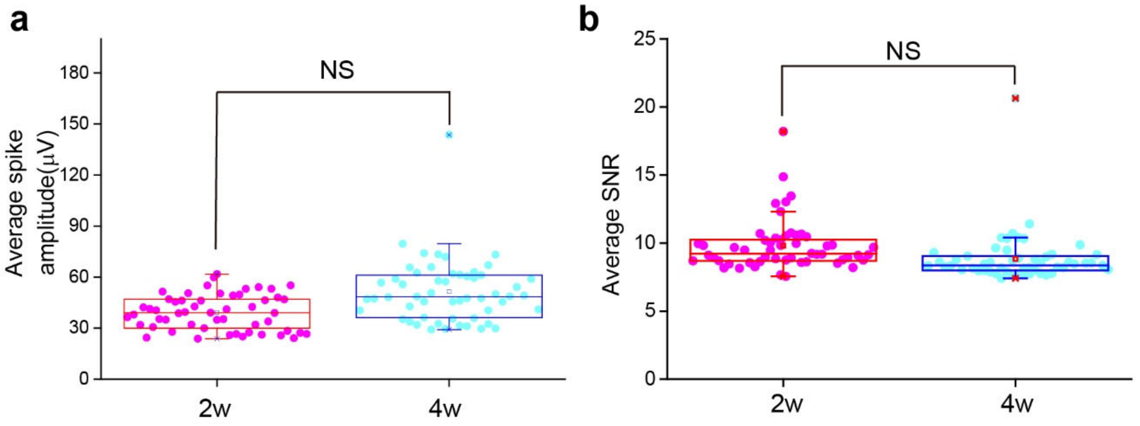


**Figure S10. Time-dependent average amplitude and SNRs. a-b**, Average spike amplitude (**a**) and SNR (**b**) from three mice at 2 weeks and 4 weeks post-injection. The total average spike amplitudes of 84 channels were ~47 μV at 2 weeks and ~50 μV at 4 weeks post-injection, and the average SNRs were ~10 at both time points. Box plots show mean (open squares), median (horizontal lines), quartiles (boxes, 25-75%), and ranges (whiskers, 1-99%). NS: not significant ($P > 0.05$, paired-sample $t$ test).

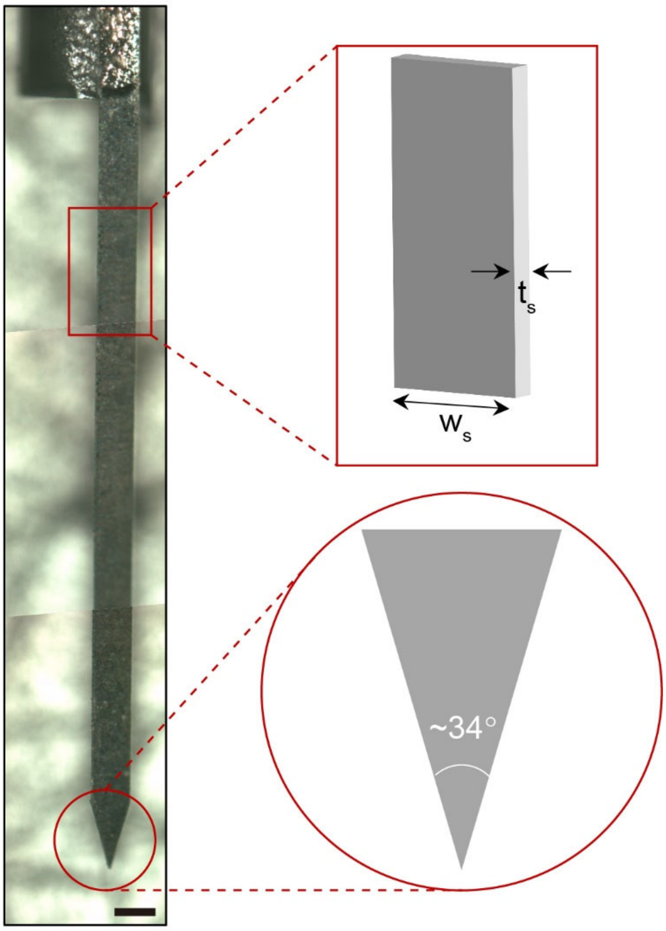


**Figure S11. Geometrical consideration of ultrathin metal insertion needle.** Stitched optical image showing the stainless metal insertion needle attached to glass capillary needle by dental cement. The thickness ($t_s$) and width ($w_s$) of metal needle are 20 μm and 200 μm, respectively. The opening angle of the metal needle is set to ~34° for penetration to the dura/brain. Scale bar, 200 μm.

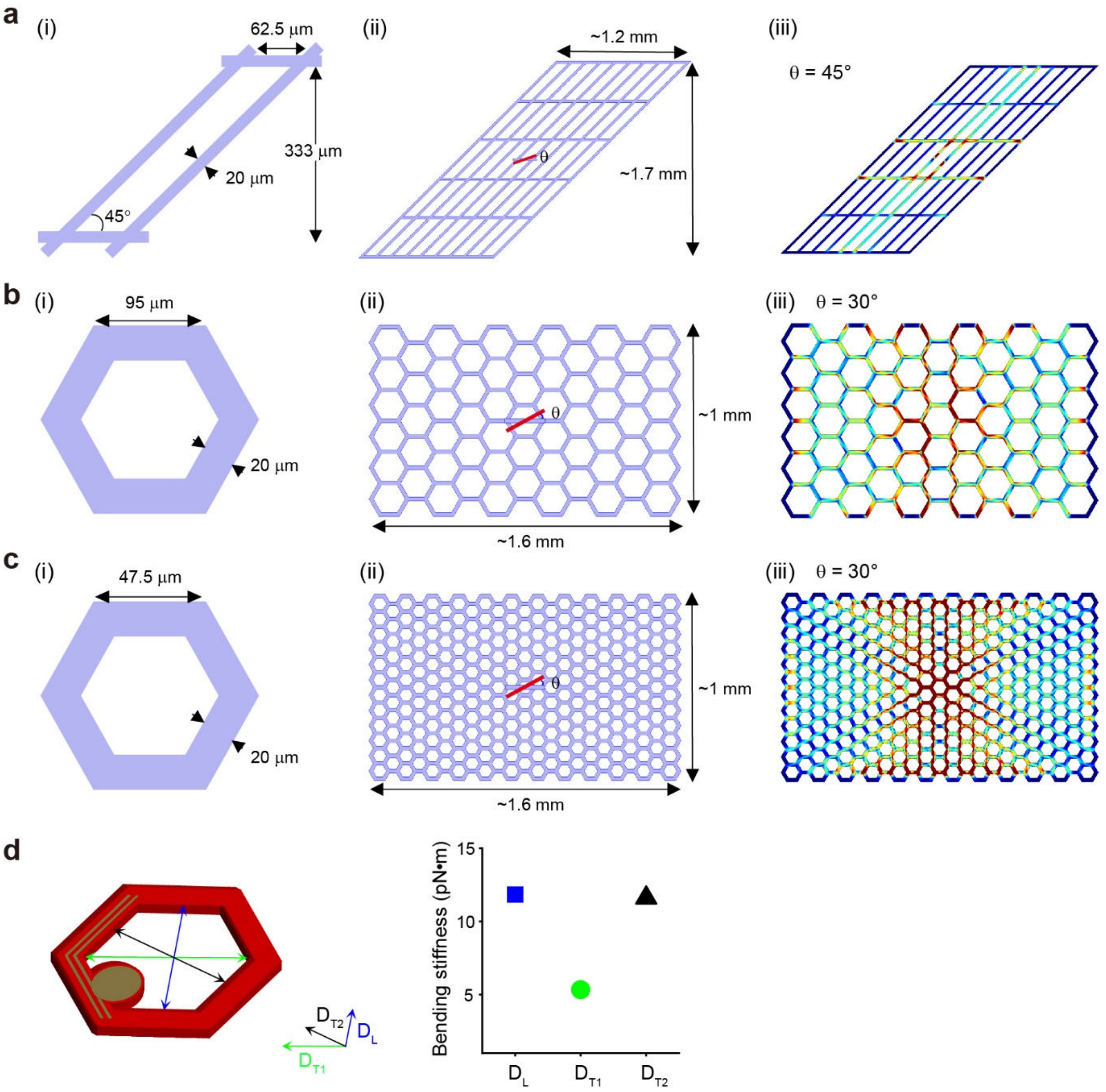


**Figure S12. Numerical simulations of von Mises stress distribution and bending stiffness. a-c,** Standard mesh (θ = 45°) (**a**), larger hexagonal shaped unit cell (θ = 30°) (**b**), and smaller hexagonal shaped unit cell (θ = 30°) (**c**) are examined for FEM simulation. Each structure is shown: (i) schematics of unit cells; (ii) full-domain structures for simulation; (iii) simulated von Mises stress distributions when the 20 μm-thickness and 200 μm-width of the rod presses the center of different designs of probes (**a-c**). For the standard mesh of **a**, the distance between transverse elements and the distance between longitudinal elements are 125 μm and 333 μm, respectively. The one-side lengths of the larger (**b**) and smaller hexagonal shaped probe (**c**) are 95 μm and 47.5 μm, respectively. The widths of SU-8 ribbons in these meshes are all 20 μm. Full domain sizes for the standard mesh, larger hexagonal shaped probe, and smaller hexagonal shaped probe are ~1.2 ×1.7 mm$^2$, ~1.6 ×1.0 mm$^2$, and ~1.6 ×1.0 mm$^2$, respectively. The pressing rod is rotated by θ from the horizontal line in the simulation. **d**, Simulated bending stiffness along the longitudinal ($D_L$, blue) and transverse directions ($D_{T1}$, green and $D_{T2}$, black) for the smaller H-mesh of **c**.

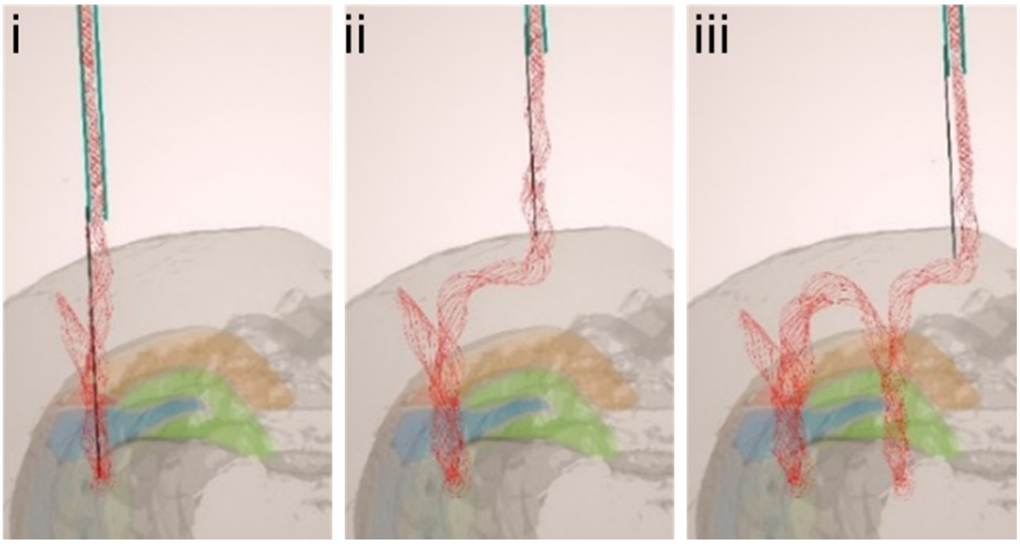


**Figure S13. Schematics and optical images of stitching hexagonal shaped probe using metal insertion needle.** A series of schematics highlighting following key steps. (i) The whole hexagonal shaped probe (red) is loaded into a large-diameter capillary needle. 1 mm-long metal needle is attached to the glass needle. (ii) The metal insertion is lifted up using motorized stereotaxic frame and moves to another target area. (iii) The processes of (i) and (ii) are repeated for two more sites implantations.

In the case of implantation using a metal insertion needle (thickness of 20 μm), the minimum separation distance between implantation sites would be ~20 μm. The maximum distance could be substantially larger than for implantation using a glass capillary needle, and ultimately, would be limited by probe fabrication (i.e., 100's mm separation between sites).

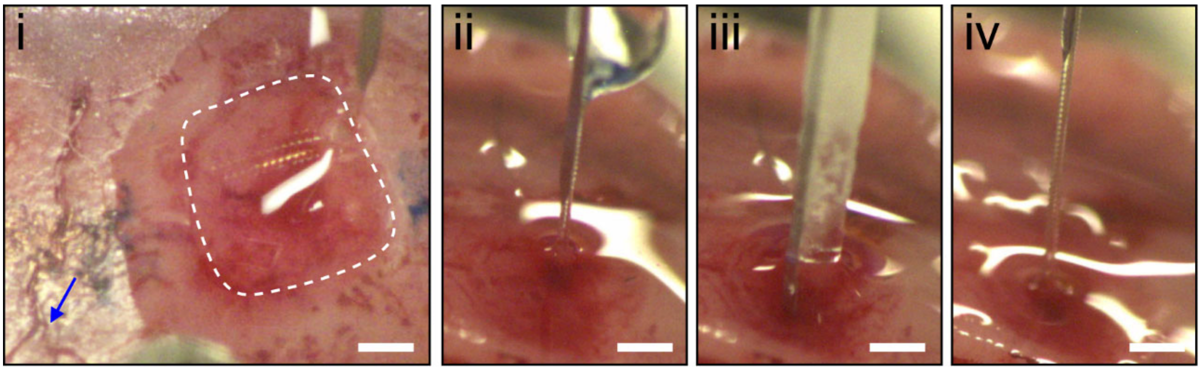


**Figure S14. Optical images of stitching hexagonal-shaped probe into mouse brain using metal insertion needle.** Optical microscope images showing the procedure of in-vivo implantation using metal insertion needle. A single hexagonal-shaped probe is implanted into the hippocampal region of a mouse brain: (i) The skull area of ~3 × 3 $mm^2$ is opened by dental drill (white dashed region in i), and the dura is removed for implantation. The blue arrow indicates the bregma. (ii)-(iv) A series of optical images showing the procedure of hexagonal shaped probe implantation using the metal insertion needle. Scale bars, 1 mm.

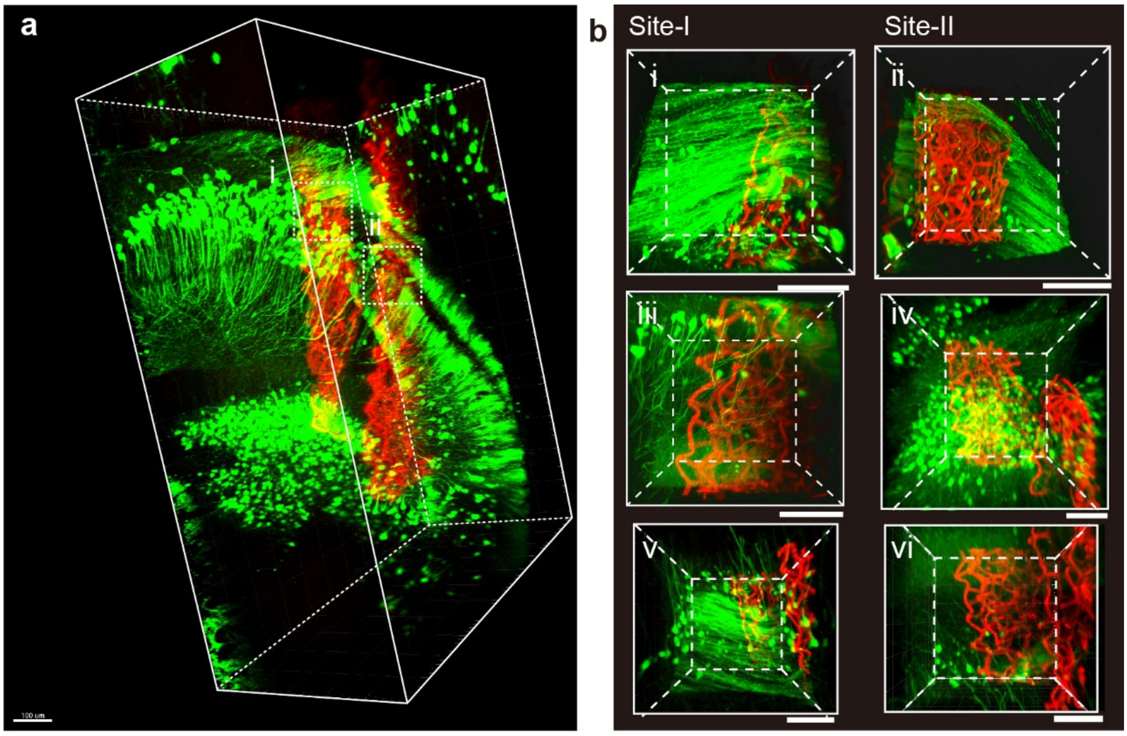


**Figure S15. High-resolution 3D mapping and analysis of hexagonal shaped probe-neural interface.** **a**, 3D full probe interfaces of YFP-H transgenic mice brain (N = 3) between neurons (green) and probe (red) at 2 weeks post-injection. Scale bars, 100 μm. **b**, High-resolution image of the volumes highlighted by white dashed boxes (i, site-I) and (ii, site-II) in **a**. Scale bars, 100 μm. (iii)-(vi) High-resolution image from the other YFP-H transgenic mouse brain at site-I (iii and v) and site-II (iv and vi). Scale bars, 100 μm.

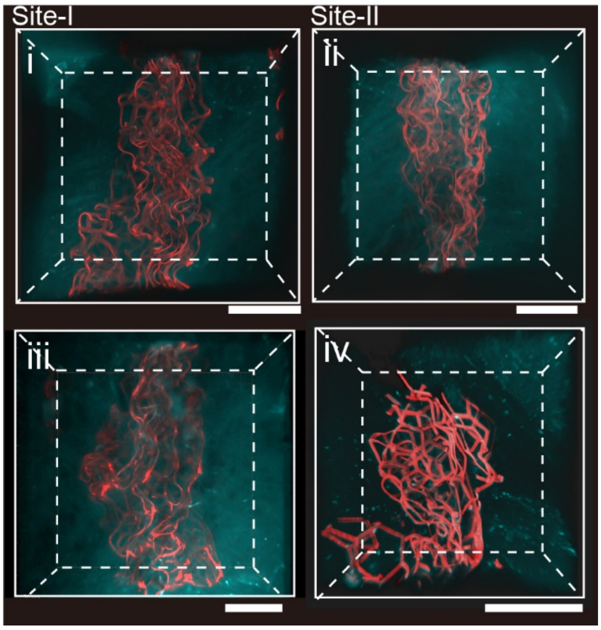


**Figure S16. High-resolution 3D mapping and analysis of hexagonal shaped probe-astrocyte interface.** High-resolution 3D reconstructed images of GFAP transgenic mice brain (N = 2) at 2 weeks post-injection. 3D images of the interface between astrocyte (cyan) and hexagonal-shaped probe (red) at site-I (i and iii) and site-II (ii and iv). Scale bars, 100 µm.

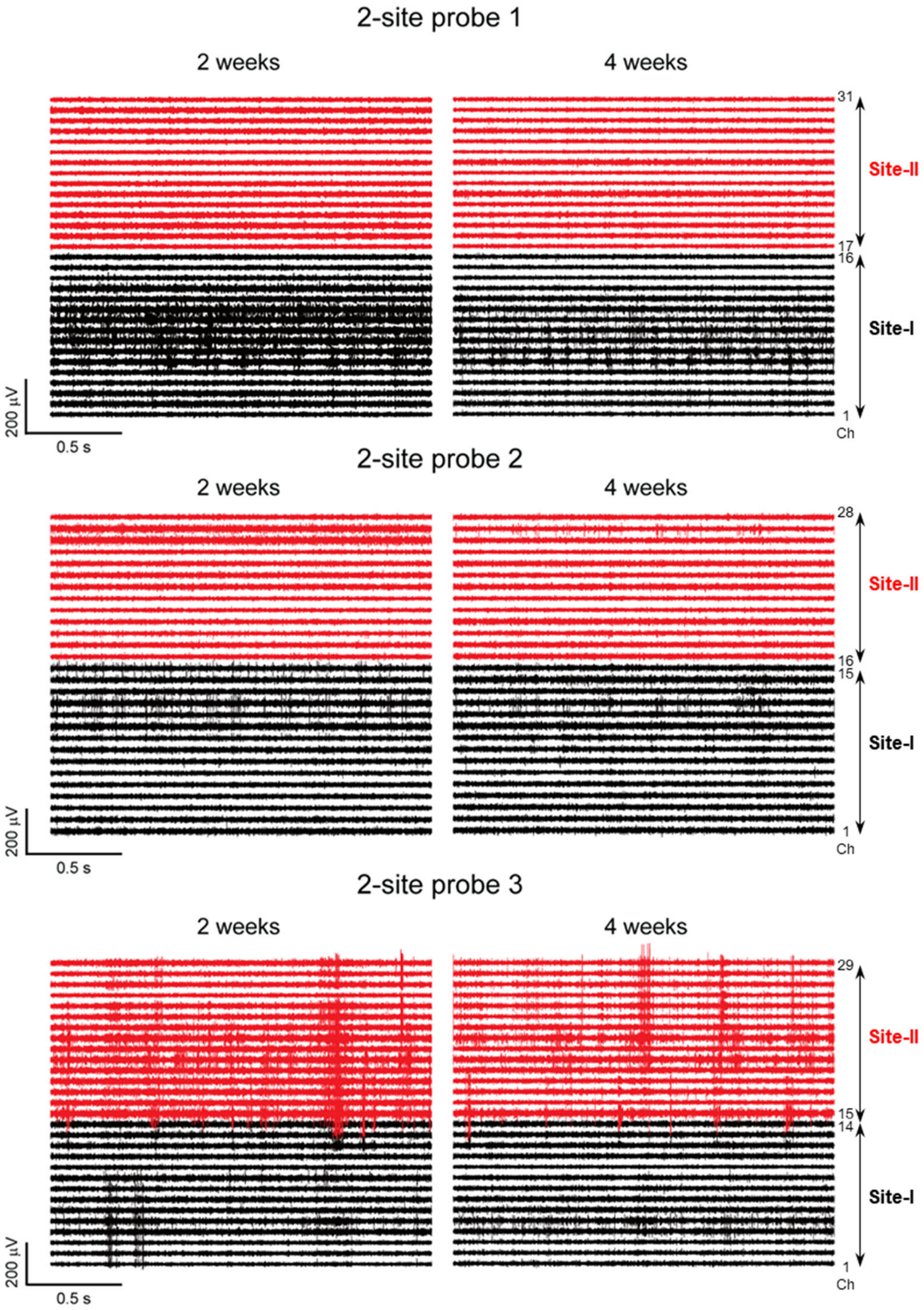


**Figure S17. Extracellular recordings by mesh probe implanted using metal insertion needle.** Band-pass (250-6000 Hz) filtered extracellular single-unit spike traces recorded after two-site implantations in three individual mice (N = 3) at 2 and 4 weeks post-injection. Black and red traces were recorded from site-I and II, respectively.

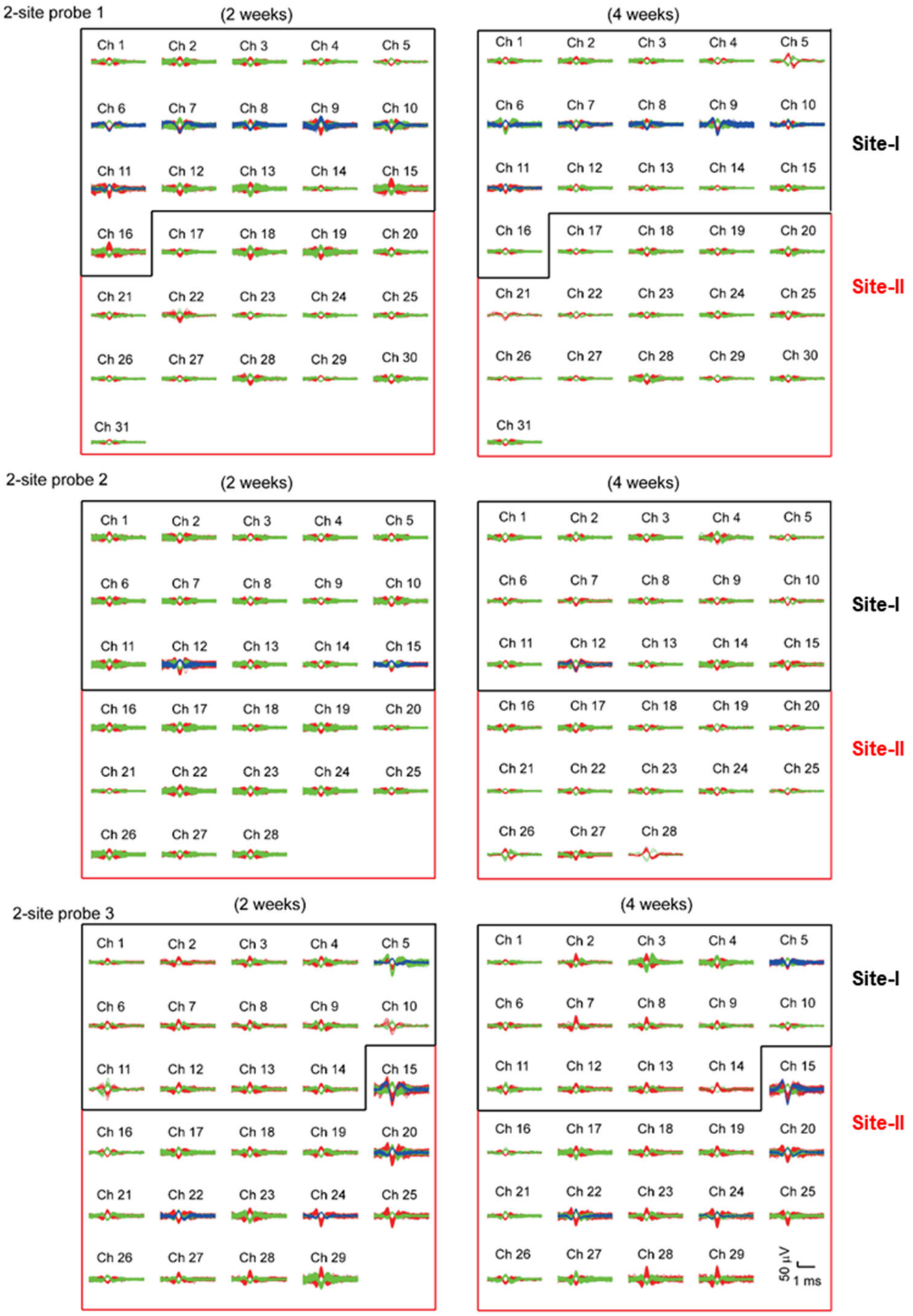


**Figure S18. Overlay of sorted and clustered spikes.** Spike sorting analysis of the recording data shown in Figure S17 (total 88 channels in three mice at 2 and 4 weeks post-injection). There are 99 (site-I) and 90 (site-II) single-unit spikes from 88 channels at 2 weeks; 98 (site-I) and 90 (site-II) single-unit spikes from 88 channels at 4 weeks.

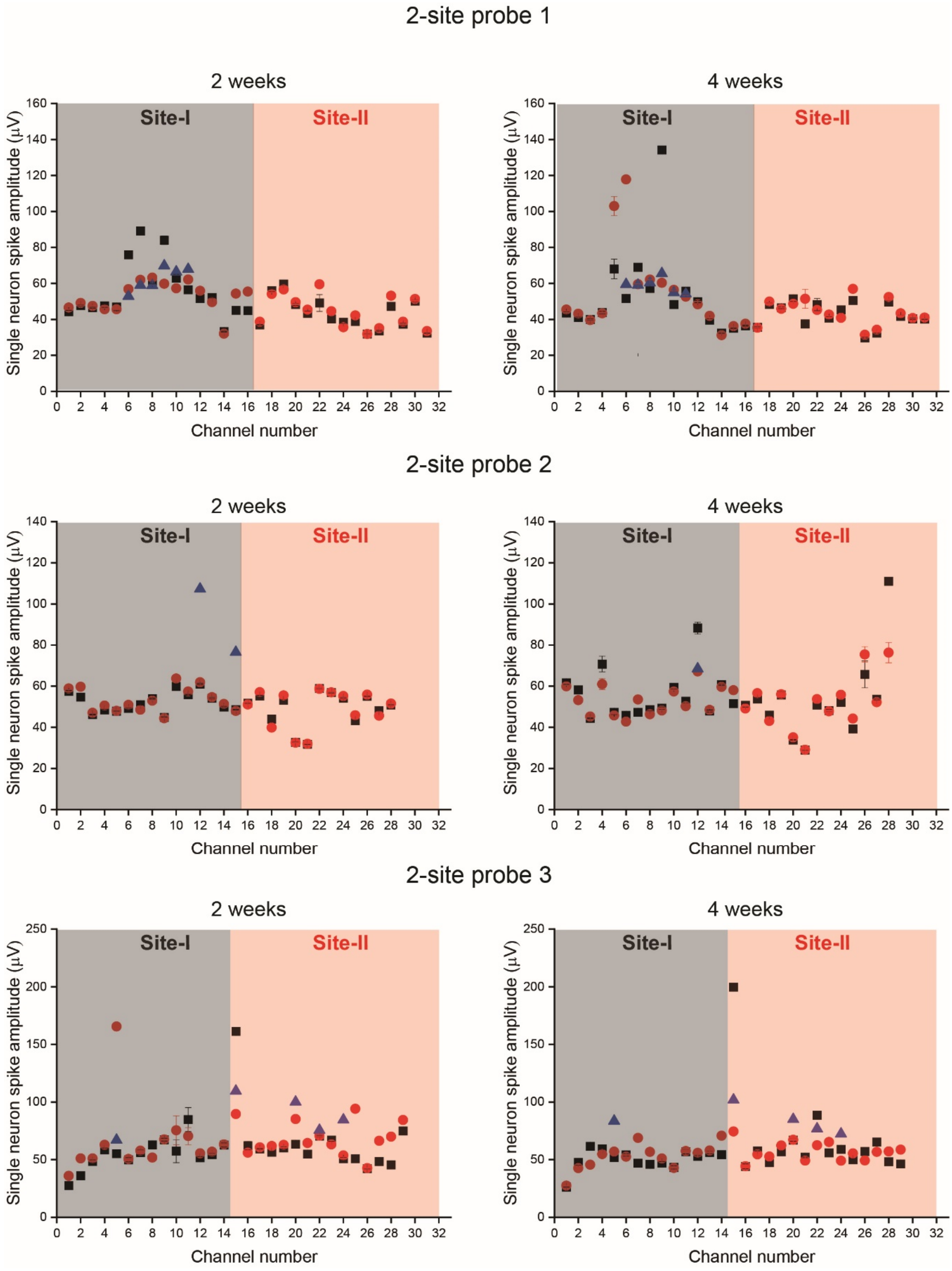


**Figure S19. Time-dependent spike amplitudes from three mice**. **a-c**, Single-unit peak-to-peak spike amplitude from three mice (N = 3) at 2 and 4 weeks. Data were obtained from Figure S17. The amplitudes of the sorted single-unit spikes at 2 weeks were ~56 μV from 45 channels and ~55 μV from 43 channels in the 1st and 2nd regions, respectively. The average spike amplitudes are ~55.9 and ~54.0 μV at 2 and 4 weeks, respectively.

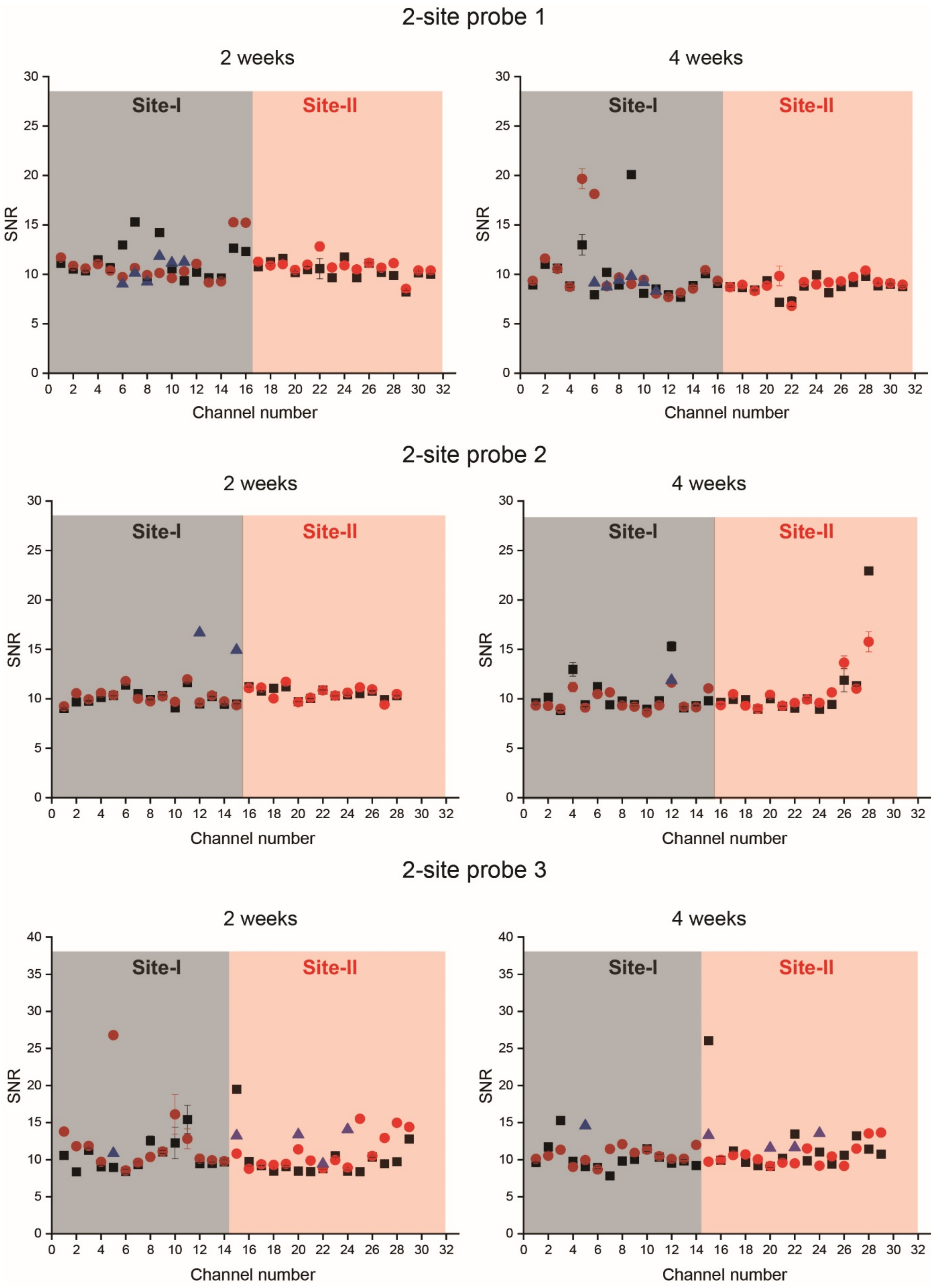


**Figure S20. Time-dependent SNRs.** Single-unit SNR from three mice (N = 3) at 2 and 4 weeks. Data were obtained from Figure S17. The average SNRs were ~10 in both site-I and site-II at 2 weeks. The average SNRs are ~10.8 and ~10.3 at 2 and 4 weeks, respectively.

### Supplementary Movie Legends

**Movie 1.** In-vitro demonstration of stitching mesh electronics in a hydrogel using a glass capillary needle. The video shows implantation into four-sites of a brain-mimicking 0.5% agarose hydrogel; the mesh design is shown in Figure S1a. The movie also shows that the flexible mesh probe was implanted in an extended conformation without crumpling at each of the four sites, and that mesh region connecting the implantation sites is on the surface of the hydrogel. The video is played at 16× real time.

**Movie 2.** In-vitro demonstration of stitching a hexagonal mesh probe using a metal insertion needle. This video shows two-site implantations into 0.5% agarose hydrogel with the design shown in Figure S1b. The whole hexagonal array probe was loaded into large capillary needle attached with metal insertion needle and implanted into hydrogel by metal insertion without the aid of liquid flow. The glass capillary needle was not inserted into hydrogel. In this case, the connector part was intentionally implanted. The video is played at 16× real time.
